# Ubiquitin-protein ligase *Ubr5* cooperates with Hedgehog signalling to promote skeletal tissue homeostasis

**DOI:** 10.1101/2020.12.04.411405

**Authors:** David Mellis, Katherine A Staines, Silvia Peluso, Ioanna Ch. Georgiou, Natalie Dora, Malgorzata Kubiak, Michela Grillo, Colin Farquharson, Elaine Kinsella, Anna Thornburn, Stuart H Ralston, Donald M Salter, Natalia A Riobo-Del Galdo, Robert E Hill, Mark Ditzel

**Author notes:** Corresponding author: (REH). These authors contributed equally to this work (David Mellis, Katherine A Staines). These authors also contributed equally to this work (Natalia A Riobo-Del Galdo, Robert E Hill, Mark Ditzel).

## Abstract

Mammalian Hedgehog (HH) signalling pathway plays an essential role in tissue homeostasis and its deregulation is linked to rheumatological disorders. UBR5 is the mammalian homologue of the E3 ubiquitin-protein ligase Hyd, a negative regulator of the Hh-pathway in *Drosophila.* To investigate a possible role of UBR5 in regulation of the musculoskeletal system through modulation of mammalian HH signaling, we created a mouse model for specific loss of *Ubr5* function in limb bud mesenchyme. Our findings revealed a role for UBR5 in maintaining cartilage homeostasis and suppressing metaplasia. *Ubr5* loss of function resulted in progressive and dramatic articular cartilage degradation, enlarged, abnormally shaped sesamoid bones and extensive heterotopic tissue metaplasia linked to calcification of tendons and ossification of synovium. Genetic suppression of smoothened *(Smo*), a key mediator of HH signalling, dramatically enhanced the *Ubr5* mutant phenotype. Analysis of HH signalling in both mouse and cell model systems revealed that loss of *Ubr5* stimulated canonical HH-signalling while also increasing PKA activity. In addition, human osteoarthritic samples revealed similar correlations between *UBR5* expression, canonical HH signalling and PKA activity markers. Our studies identified a crucial function for the *Ubr5* gene in the maintenance of skeletal tissue homeostasis and an unexpected mode of regulation of the HH signalling pathway.

**Author Summary:** Ubiquitin ligases modify proteins post-translationally which is essential for a variety of cellular processes. UBR5 is an E3 ubiquitin ligase and in *Drosophila* is a regulator of Hedgehog signaling. In mammals, the Hedgehog (HH) signalling pathway, among many other roles, plays an essential role in tissue maintenance, a process called homeostasis. A murine genetic system was developed to specifically eliminate UBR5 function from embryonic limb tissue that subsequently forms bone and connective tissue (ligaments and tendons). This approach revealed that UBR5 operates as a potent suppressor of excessive growth of normal cartilage and bone and prevents formation of bone in ectopic sites in connective tissue near the knees and ankle joints. In contrast to abnormal growth, UBR5 inhibits degradation of the articular cartilage that cushions the knee joint leading to extensive exposure of underlying bone. Furthermore, Ubr5 interacts with smoothened, a component of the HH pathway, identifying UBR5 as a regulator of mammalian HH signaling in the postnatal musculoskeletal system. In summary, this work shows that UBR5 interacts with the HH pathway to regulate skeletal homeostasis in and around joints of the legs and identifies targets that may be harnessed for biomedical engineering and clinical applications.

## Introduction

Ubiquitin ligases target proteins for ubiquitination which can modulate protein function by regulating protein degradaton, protein–protein interactions, and protein localization [1–4], and thus, provide important post-translational mechanisms essential for a variety of cellular processes. The *Drosophila* homologue of the mammalian *Ubiquitin Protein Ligase E3 Component N-Recognin* 5 (UBR5), designated as *hyperplastic discs* (Hyd), was originally identified as a Drosophila tumor suppressor protein [5–7] and regulator of Hedgehog (HH) signalling [6]. Physical and genetic interactions with established components of the HH signalling pathway [7, 8] strengthened Hyd’s role as a regulator of HH signalling. We previously addressed a possible conserved role for UBR5 in HH-mediated processes in mice [9]. Although no overt effects were seen in patterning of the developing limb bud in mouse embryogenesis; here, we show that the coordinated action of Ubr5 with HH signalling is crucial to maintain skeletal tissue homeostasis associated with the appendicular skeleton postnatally and in adult mice.

HH signalling regulates cell processes that are critical for skeletal tissue development, growth and homeostasis [10]. Two HH ligands, Sonic- and Indian-Hedgehog (SHH and IHH, respectively) are widely expressed and function as extracellular signalling molecules that bind to cells expressing HH receptors such as patched-1 (PTCH1). Binding to PTCH1 results in de-repression of the G protein-coupled receptor, smoothened (SMO), and activation of SMO-associated canonical and non-canonical signalling pathways [11–13]. Activation of the SMO-associated canonical pathway results in stimulation of GLI-mediated transcription and expression of crucial target genes [7]. Activation of the recently identified SMO-associated non-canonical pathway relies on SMO’s GPCR activity [14, 15] and results in inhibitory heterotrimeric G protein-mediated inhibition of adenylate cyclase and a concomitant reduction in cyclic AMP (cAMP) levels [14, 16, 17]. Although not yet experimentally addressed, non-canonical signalling may also contribute to many of the well-described roles for canonical HH signalling in normal skeletal formation, maturation and maintenance [10, 18].

At birth, IHH is the ligand that drives HH signalling within the growing limbs. Expression of *Ihh* is localized to a zone of postmitotic, prehypertrophic chondrocytes immediately adjacent to the zone of proliferating chondrocytes [18–20] and is essential for endochondral ossification but also induces osteoblast differentiation in the perichondrium [21]. Dysregulation of this signalling pathway is detrimental to musculoskeletal tissue homeostasis [22, 23]. Notably, studies have shown that increased HH signalling can drive pathological ectopic cartilage and bone formation in soft tissues [10] through the process of heterotopic chondrogenesis and heterotopic ossification (HO) [24]. Upregulation of HH signalling is believed to contribute to the rare disorder, progressive osseous heteroplasia (POH), which includes in its phenotypic spectrum soft tissue ossification. POH is caused by loss-of-function of *GNAS,* a G protein alpha subunit and activator of adenylate cyclase. A murine model of POH demonstrated that increased HH signalling as a consequence of *GNAS* loss-of-function in mesenchymal limb progenitor cells drove heterotopic ossification [25]. Similarly, synovial chondromatosis, a disease resulting in ossification of synovial tissue is associated with increased canonical HH signalling [26]. However, in contrast with cartilage and bone gain, elevated HH signalling is also associated with the cartilage degradation and loss [27, 28]. Hence, appropriate HH signalling is normally involved in the suppression of ectopic, and genesis and maintenance of normtopic, cartilage and bone.

Here, we show that the loss of *Ubr5* function in *Ubr5^mt^* mice resulted in diverse musculoskeletal defects including spontaneous, progressive and tissue-specific patterns of ectopic chondrogenesis and ossification as well as articular cartilage degeneration and shedding. Surprisingly, reducing SMO function in UBR5-deficient mice led to a dramatic reduction in the age of onset and increased severity of the *Ubr5^mt^* phenotype. These observations challenge the existing dogma by highlighting an important role for *Smo,* in the absence of UBR5, in suppressing, rather than promoting, ectopic chondrogenesis, tissue calcification/ossification and articular cartilage damage. We, therefore, reveal a previously unknown physiological role for *Ubr5* and highlight its genetic interaction with *Smo* in regulating cellular and tissue-homeostasis. These findings may influence current therapeutic approaches modulating HH signalling for the treatment of osteoarthritis and heterotopic ossification.

## Results

### Loss of *Ubr5* function causes skeletal heterotopias at 6 months

To overcome the embryonic lethality associated with germline mutant animals [29], we combined a *Ubr5* conditional loss-of-function gene trap *(Ubr5^gt^)* [9] with *Prx1-Cre* [30] *(Prx1-Cre;Ubr5^gt/gt^* animals henceforth, referred to as *Ubr5^mt^)* to ensure that adult tissues derived from early limb bud mesenchyme, predominantly bone and connective tissue, were *Ubr5* deficient. Since the HH pathway affects embryonic limb patterning and bone growth, the *Ubr5* deficient fetuses (at E15.5) were initially examined and bones and joints appeared to develop normally [9]. However, the HH pathway continues to function in postnatal bone growth and homeostasis [10] and thus, at approximately 6 months of age, we noticed that mice began to display defects in locomotion. Control animals normally remained supported by their hindlimbs (‘sprung’), whereas, *Ubr5^mt^* animals rested their posteriors directly upon the floor (‘squat’) (S1 Fig A-C). Considering the tissue targeted by the conditional mutation, the observed phenotype indicated a potential musculoskeletal system defect which prompted the examination of hindleg bone and joint structures.

At 6 months of age, X-ray imaging revealed that *Ubr5^mt^* animals exhibited abnormally shaped and/or ectopic signals around knee and ankle joints (S1 Fig D-I). 3D micro-computed tomography (μCT) revealed that, whereas *Prx1-Cre* control joints appeared normal with no evidence of ectopic structures (Fig 1A), the knees and ankles of all *Ubr5^mt^* mice (n=10) exhibited isolated ectopic signals clearly separated from the adjacent femoral condyles and tibia (Fig 1B). Surface rendering of the μCT scans demonstrated that the array of knee-associated sesamoid bones (patella and fabella) and calcified menisci (Fig 1 C,D) were abnormal. *Ubr5^mt^* knees presented with large ectopic structures on all four faces of the knee joint, as well as enlarged and irregularly shaped fabella and patella sesamoid bones (Fig 1D). In addition, the *Ubr5^mt^* animals exhibited multiple ectopic signals around the ankle joint (Fig 1 E-G), with the most striking one appearing consistently on the dorsal side running parallel to the long axis of the tibia (Fig 1 F, open arrows) associated with the Achilles tendon (AT). This ectopic signal remained isolated from the calcaneus and tibia. Other ectopic structures included two ectopic U-shaped signals on the ventral and lateral sides of the tibia (Fig 1 G).

**Figure 1.**
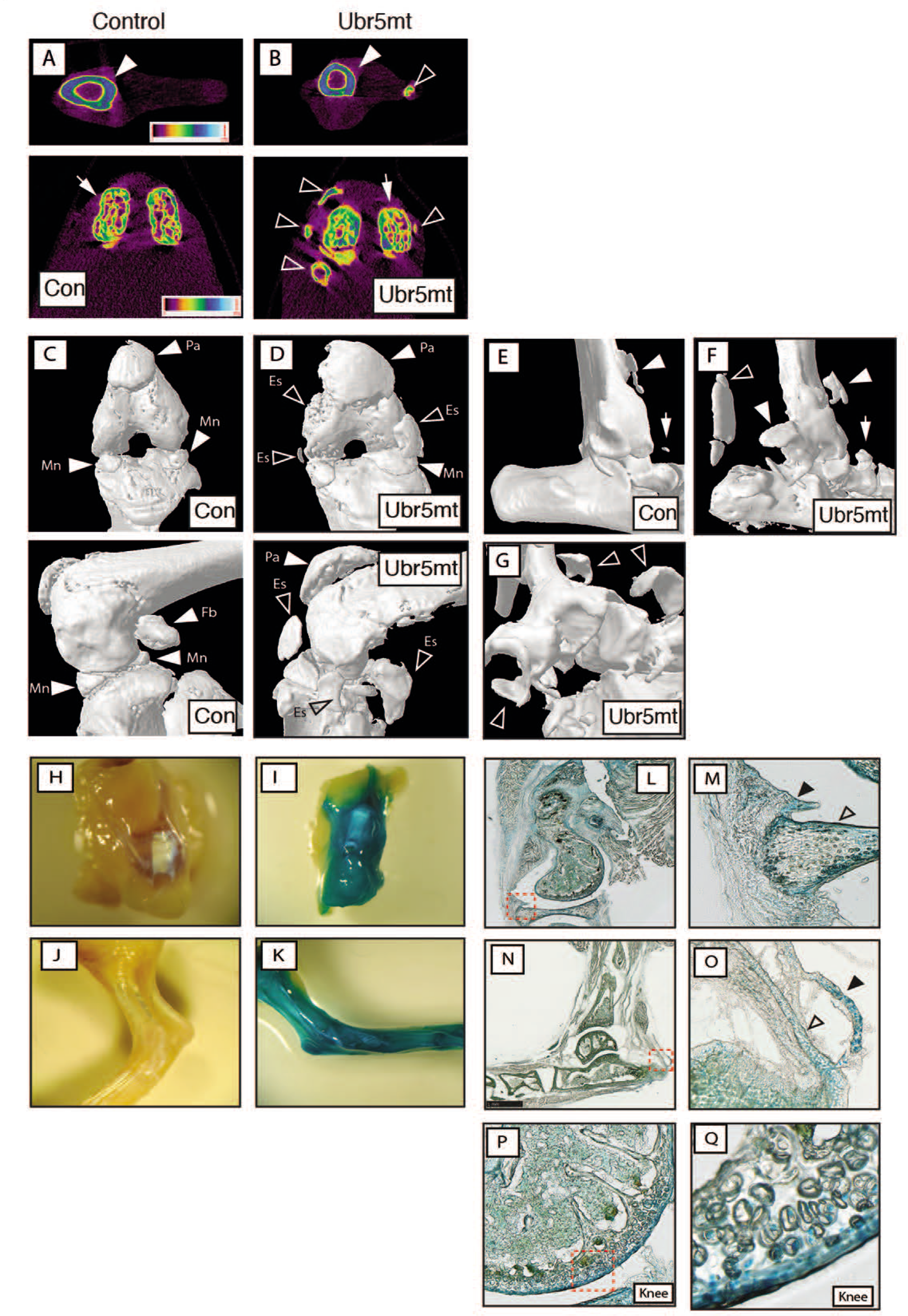
*Ubr5^mt^* animals exhibit multiple ectopic structures around the knee and ankle joints. 6-month old control and *Ubr5^mt^* animals were analysed by μCT. (A, B) Color-coded density maps revealed presence of ectopic, non-uniform density structures (open arrowheads) around the tibia (closed arrowheads) and femoral condyles (arrows). (C, D) Knee joints in ventral (upper panels) or medial (lower panels) aspect. Closed arrowheads indicate normal structures: the patella (Pa), menisci (Mn) and fabella (Fb). Open arrowheads indicate ectopic signals (Es) present in the *Ubr5^mt^* knee joint. (E) Control ankle joints exhibit a signal extending from the ventral face of the tibia (closed arrowhead) and a small structure presumed to be a sesamoid bone (arrow). (F) Multiple ectopic signals were present around the *Ubr5^mt^* ankle joint (arrowheads), including a large dorsally located and well-isolated structure in the location of the AT (open arrowhead). (G) Higher magnification of multiple ectopic signals (open arrowheads). (H-Q) 20-week-old *Prx1-Cre* control and *Ubr5^mt^* ankle and knee joints were stained for β-gal activity. Whole mount knee (H,I) and ankle (J,K) are shown with subsequent sagital sections for the knee (L) and the boxed area magnified in (M) shows the outer layer of the menisci (open arrowhead), and the adjacent synovium (closed arrowhead) stained positive for β-gal expression. Sagital sections for the ankle are shown in (N) and magnified in (O) showing staining of the AT and superficial digital flexor tendon (open and closed arrowheads, respectively). (P,Q) Expression of β-gal in the articular cartilage of the knee and the box in P shown at higher magnification in Q.

Following recombination of the *Ubr5^gt^* gene-trap construct, *lacZ* is expressed under the influence of *Ubr5* gene regulators enabling the analysis of the postnatal tissues expressing *Ubr5* Previously [9], we showed that β-gal activity was restricted to the limb mesenchyme at embryonic stages. Analysis of *lacZ* expression in 20 week-old mice control and *Ubr5^mt^* knee (Fig 1 H, I) and ankle (Fig 1 J, K) joints revealed strong β-gal activity in tissue derived from this embryonic mesenchyme. Expression occurred around the periphery of the menisci and synovium (Fig 1 L, M). The ankle also revealed β-gal activity within the AT and superficial digital flexor tendon and in a large ectopic structure within the AT midbody (Fig 1 N, O). In addition, expression was detected within the upper layer chondrocytes of the femoral and tibial articular cartilage (AC) (Fig P, Q). Thus, the tissues that exhibit Cre-mediated expression of the *lacZ* gene are affected in the mutant phenotype.

### *Ubr5^mt^*-associated ectopic structures exhibit chondrogenesis and calcification

The morphology of these ectopic structures was further investigated to determine the cellular composition and possible derivation of these ectopias. As shown by μCT, both knee (Fig. 2 A,_B) and ankle (Fig. 2 C,_D) ectopic structures harbored different X-ray densities and internal structures indicative of bone. This was confirmed in the ankle joint by von Kossa staining, in which large ectopic staining was observed in the AT (Fig. 2E, F) and in the superficial digital flexor tendon (Fig. 2E, white arrowhead). Subsequent histological analysis of the AT revealed, in *Prx1-Cre* controls, the expected ordered stacking of tenocytes along the anterior-posterior axis of the tendon (Fig. 2G) and an absence of toluidine blue staining associated with proteoglycans (Fig. 2H). In contrast, regions of the *Ubr5^mt^* Achilles tendon were devoid of tenocytes, which were replaced by long columns of proteoglycan-expressing hypertrophic chondrocytes (Fig. 2I, J). The combination of the distinctive cell morphology and toluidine blue-staining pattern suggested that ectopic chondrocytes and their associated extracellular matrix were present in *Ubr5^mt^* tendons.

**Figure 2.**
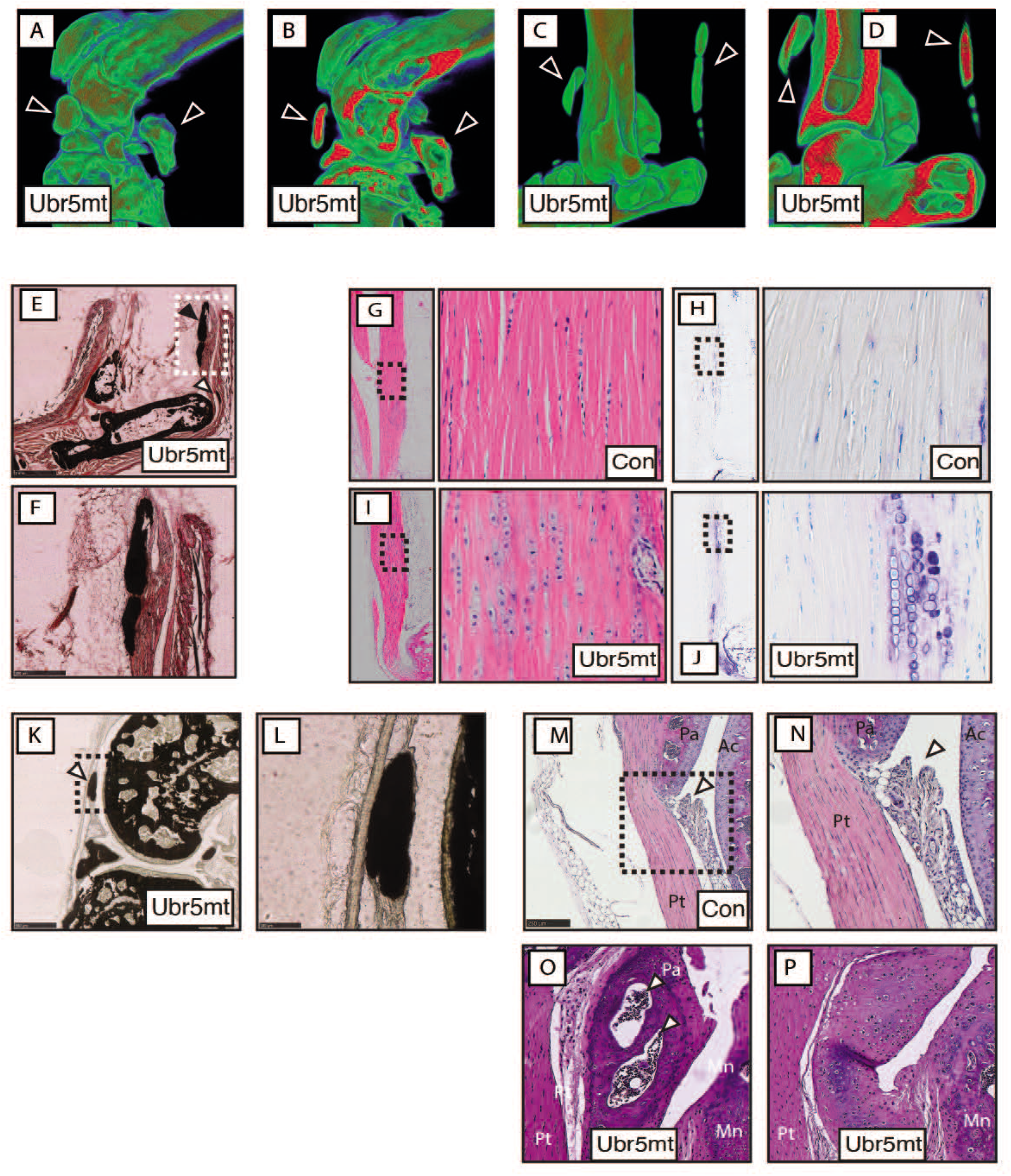
*Ubr5^mt^* limbs exhibit ectopic chondrogenesis, cartilage formation and calcification and ossification. (A-D) Colour-coded X-ray density maps of volume rendered *Ubr5^mt^* knee (A,B) and ankle (C,D) joints. (B,D) show cross-sections through the joint to reveal the internal structure and density. Arrowheads indicate ectopic structures. Low density = blue and High density = red. Sagittal sections from (E-L) 20-week-old or (M-P) 24-week-old animals are shown. (E,F) Von Kossa staining of *Ubr5^mt^* ankle joints revealed ectopic signals in AT (arrowheads). The dashed boxed region is enlarged in (F) and shows the shape and location of the ectopic structure on the deep face of the AT. (G,I) H&E and (H,J) toluidine blue staining of the midbody of Achilles tendons. The left panel of each pair shows a low magnification image of the tendon. A higher magnification of the boxed region is shown in the right panel. (G,H) Control tendons showed the expected columns of tenocytes and very little toluidine blue staining. (I,J) *Ubr5^mt^* tendons harbour chondrocytes that coincide with regions of toluidine blue staining. (K) Von Kossa staining of *Ubr5^mt^* knee joints revealed ectopic signals in the synovium (arrowhead). (L) shows an enlarged image of the ectopic structure lying within the synovium and under the patellar tendon. (M,N) Image of control synovium (arrowhead) located underneath the patella (Pa) and patellar tendon (PT) and adjacent to the tibial articular cartilage (AC). The boxed area in (M) is enlarged in (N). (O-P) *Ubr5^mt^* synovium harbours ectopic tissue. (O) The synovium harbours a bone-like structure (arrowhead). (P) In other regions, the synovium abutting the patella appeared thickened but not ossified showing cartilage harbouring chondrocytes. Pt = patellar tendon; Pa = patella; Mn = meniscus

To address the presence of ectopic calcium deposition, we used Von Kossa staining of *Ubr5^mt^* knee joints that revealed positive stained structures within the synovium deep to the patellar tendon (Fig. 2K, L). Histological analysis of *Prx1-Cre* control knee joints revealed a synoviocyte-rich intimal layer of the synovium (Fig. 2M, N), whereas *Ubr5^mt^* knee joints exhibited bone- (Fig. 2O) and cartilage-like (Fig. 2P) ectopic structures. Thus, we observed a phenotype consisting of ectopic chondrogenesis, calcification and ossification (hereafter, referred as ECCO) of the synovium and tendons in *Ubr5^mt^* tissues. We concluded that *Ubr5* normally prevents spontaneous ectopic formation of chondrocytes in tissues and calcification and/or ossification in cartilage.

### Loss of *Ubr5* function causes articular cartilage degradation

μCT analysis of 6-month old control (Fig. 3A-C) and *Ubr5^mt^* (Fig. 3D-F) knee joints revealed significantly increased volume of high subchondral bone density in the mutant (quantified in Fig 3G). Histological assessment showed a dramatic loss of articular cartilage (AC) from the lateral tibial and femoral surfaces of all *Ubr5^mt^* knee joints assessed (Fig 3H, I, K); a condition not detected in any control mice at this stage. Further examination of the exposed subchondral bone in these *Ubr5^mt^* mice revealed abnormal intermixed bone and cartilage within this region (Fig 3J). Hence, the hindlegs at 24 weeks present a diverse range of cartilaginous defects including metaplastic conversion of connective tissue associated with the knee and ankle (as described above) whereas, the AC undergoes severe degradation causing exposure of the subchondral bone at the joint surface.

**Figure 3.**
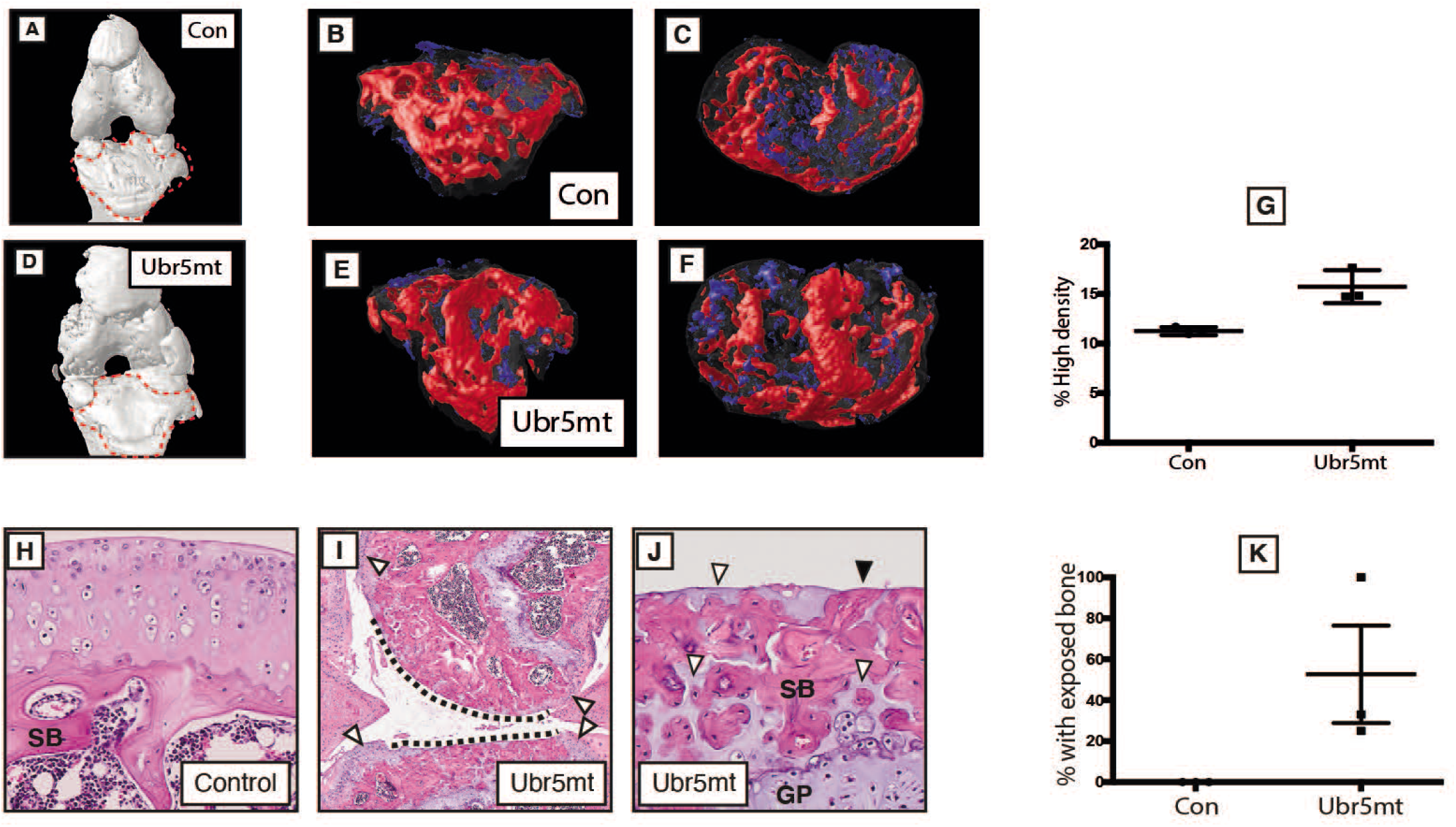
*Ubr5^mt^* animals show subchondral bone defects and AC cellular and extracellular abnormalities. (A, D) Surface rendered μCT-based 3D models images of knee joints of *Prx1-Cre* (Control) (A) and *Ubr5^mt^* (D). Volume rendered 3D models of 26-week-old tibial subchondral bone; (B, E) ventral, and (C, F) anterior views. Pan-density volume shown in grey, high-density in red and low-density in blue. (G) Graph of percentage of high-density signal volume as a percentage of total subchondral bone volume. s.e.m indicated. n = three biological replicates per genotype. t-test. p = 0.0103. H&E-based histological analysis of (H-J) 26-week-old *Prx1-Cre* (Control) and *Ubr5^mt^* tissues. (H) 26-week-old control tibial AC was normal and (I) *Ubr5^mt^* AC exhibited regions that lacked AC and exposed subchondral bone (dashed lines). Peripheral regions retained some AC (white arrowheads). (J) Closer examination revealed exposed subchondral bone (black arrowheads) and intercalated cartilage (white arrowheads). (K) Graph of % sections with exposed bone in 26-week-old tibial AC reveal a significant increase in *Ubr5^mt^* AC. Fishers exact test on pooled slide counts, p=0.0075.

### Ubr5 deficiency results in a postnatal, progressive phenotype

To establish the approximate age at which this striking ECCO phenotype is initially detectable, a timed series of *in vivo* μCT scans on ageing, live animals was followed. *Ubr5^mt^* animals at 3-weeks of age revealed no marked difference in knee or ankle joints (S2 Fig A-D), suggesting that the ectopic structures did not form during fetal development but rather formed postnatally. Between 6 and 12 weeks of age, the ectopic structures began to emerge (Fig 4A, B), initially on the ventral side of the tibia. Dorsally located ectopic signals associated with the Achilles’ tendon emerged by 16 weeks of age (Fig 4C) and all ectopic structures were enlarged by 24 weeks of age (Fig 4D). These data suggest that *Ubr5* deficiency led to enhanced, progressive chondrogenesis and osteogenesis in the connective tissue.

**Figure 4.**
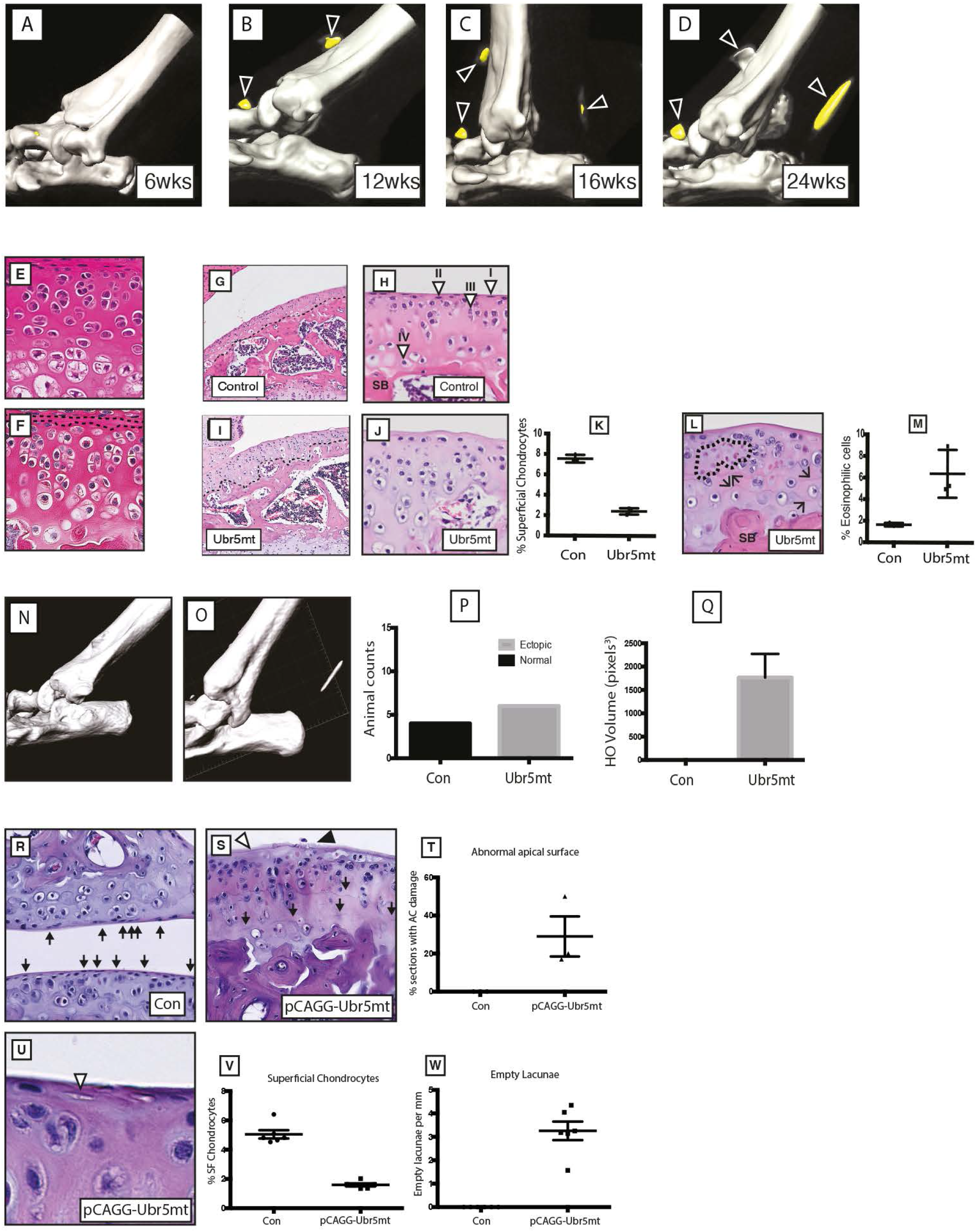
Ubr5mt mice exhibit degenerative, age related defects. (A-D) Consecutive uCT scans of a live *Ubr5^mt^* animals at the indicated ages. *Ubr5^mt^* ankles form (i) ventral ectopic signals (arrowheads) around (B) 12 weeks of age and (ii) dorsal ectopic signals around (C) 16 weeks of age that (D) increased in size over time. Three-week-old knee joints, stained with H&E, of *Prx1-Cre* (control) (E) and *Ubr5^mt^* (F). Black dashed lines in (F) demarcate the *Ubr5^mt^* apical acellular region. (G-M) Six-week-old (G,H) *Prx1-Cre* (control) and (I,J,L) *Ubr5^mt^* proximal tibial AC were analysed. (G,H) Control AC revealed the expected chondrocyte profile along the apical to basal axis, namely (I) superficial chondrocytes lining the apical surface; (II) non-hypertrophic rounded/oblong nuclei chondrocytes; (III) larger pre-hypertrophic-like chondrocytes within the central zone; and (IV) large hypertrophic chondrocytes located near the border with the underlying subchondral bone. SB = subchondral bone. (I) *Ubr5^mt^* tibial AC revealed abnormal chondrocytes and an acellular apical layer lacking superficial chondrocytes. (K) Graph of the percentage of superficial chondrocytes in six-week-old tibial AC. N = three biological replicates of each genotype. Mean and s.e.m indicated. Chi square test on pooled cell counts. p = <0.0001. (L) *Ubr5mt* tibial AC exhibited clusters of eosin positive chondrocytes (dashed lines) and multiple tidemarks (arrows). (M) Graph of percentage of eosinophilic chondrocytes in six-week-old AC. Chi square test on pooled cell counts. N = three biological replicates of each genotype. p = <0.001. (N-Q) Postnatal *pCAGG-Cre*-mediated recombination of the Ubr5^gt^ construct (O), but not *pCAGG-Cre* expression alone (N), resulted in X-ray dense ectopic signals forming in the AT region. (P) Counts of animals exhibiting ankle-associated ectopic signals, scored for the absence (Normal) or presence of ectopic signals (Ectopic), n = >4 for each genotype. Fisher’s exact test, p value = 0.0048. (Q) Volumetric measurement of ectopic signals from each animal n= 4. Unpaired t test, p = 0.0079. Standard error indicated. Control AC (R) exhibited superficial chondrocytes (arrows). (S) *pCAGG-Ubr5^mt^* AC exhibited an acellular apical layer (arrowheads), multiple tidemarks (arrows), and surface damage (black arrowhead). (T) Graph of percentage of sections with acellular regions and AC damage. Mean and s.e.m indicated. n = three biological replicates. Three slides analysed from each animal. Average plotted. Fishers exact on pooled section counts. p = 0.0434. (U) *pCAGG-Ubr5^mt^* AC also exhibited a reduction in superficial chondrocytes and an increase in empty apically-located lacunae (arrowheads). Graph of (V) superficial chondrocytes and (W) empty lacunae expressed as number per mm of AC. n = three biological replicates. Analysis of two sections per animal. Individual slide values plotted. Mean and s.d indicated. Fishers exact test on pooled counts. (E, F) p = <0.0001.

These metaplastic conversions within the connective tissue supporting the knee and ankle, however, contrast with the changes demonstrated in the AC which manifests as a degenerative phenotype. To investigate the timing of AC degradation, we examined mice at 3 and 6 weeks. No gross structural disruption of the AC in the *Ubr5^mt^* animals at 3-weeks of age was detected (Fig 4E, F). By 6-weeks of age, *Ubr5^mt^* articular cartilage exhibited an irregular osteochondral interface (Fig 4G, I), clusters of large, hypertrohic-like chondrocytes (Fig 4 H, J) and a reduction in the number of superficial chondrocytes (Fig 4K). *Ubr5^mt^* articular cartilage also exhibited multiple tidemarks and regions of strongly eosin positive nuclei indicative of necrosis (Fig 4L, M) that were absent in controls. The loss of Ubr5 function, therefore, resulted in early cellular and extracellular AC abnormalities prior to the progressive AC degradation, increased subchondral bone density and exposure of subchondral bone detected in 6-month old animals.

Despite loss of UBR5 in early limb mesenchyme, these data indicated that the ectopic structures arose postnatally and subsequently progressed with age. To directly address if postnatal UBR5 function was required to suppress ECCO and the degradation of the AC, we utilised a mouse line carrying a tamoxifen-inducible, conditional Cre, *pCAGG-CreERT2* [30]. Control *pCAGG-CreERT2 (pCAGG-Con)* or *pCAGG-CreERT2;Ubr5^gt/gt^* (*pCAGG-Ubr5^mt^*) animals were treated with tamoxifen (administered on two consecutive days) at six weeks of age. Staining for β-gal activity, although more broadly distributed, confirmed tamoxifen-mediated recombination of the *Ubr5^mt^* gene trap and its associated β-gal expression in tissues that included muscles and tendons (S2 Fig E, F), and within the midbody ectopia at the AT(S2 Fig G). μCT analysis at 8 weeks revealed that tamoxifen-treated control animals exhibited no ectopic signals (Fig 4N), whereas *pCAGG-Ubr5^mt^* animals exhibited Achilles’ tendon -associated ectopic signals (Fig 4O). Scoring (Fig 4P) and heterotopic ossification (HO) volumetric analysis (Fig. 4Q) confirmed that only tamoxifen-treated *pCAGG-Ubr5^mt^* animals exhibited ectopic signals. Comparison of 12 week control to treated *pCAGG-Ubr5^mt^* (Fig 4 R, S) knees revealed *Ubr5mt*-associated apical acellular layer (Fig 4S, T), damage to the apical surface, multiple tidemarks, reduced superficial zone chondrocytes (Fig 4V) and increased numbers of empty lacunae (Fig. 4U, W). We concluded that postnatal *Ubr5* function was both necessary and sufficient to maintain AC homeostasis and prevent ECCO.

### Inhibition of *Smo* promotes *Ubr5^mt^*-associated ECCO and enhances *Ubr5^mt^*-mediated AC degradation

As UBR5/HYD regulates HH signalling in Drosophila [7, 8], we next used a genetic approach to address whether aberrant HH signaling contributed to the *Ubr5^mt^* ECCO and AC phenotypes. The *Smo* gene encodes a core membrane component, regulated by the HH receptor PTCH1, that initiates the downstream signalling cascade leading to GLI-dependent transcription (canonical signalling) or Gi protein-dependent events that are tissue specific (non-canonical signalling). We reasoned that reduction in *Smo* expression levels would sensitize the HH pathway; thus, heterozygosity for a *Smo* loss of function allele (*Smo^LoF^*) [31] was used in a cross to *Ubr5^mt^* to create *Prx1-Cre;Ubr5^gt/gt^;Smo^LoF/+^* animals *(Ubr5^mt^+Smo^LoF^).*

In contrast to our expectations, μCT analysis of 12-week *Ubr5^mt^+Smo^LoF^* mice exhibited significantly more severe defects than those of age-matched *Ubr5^mt^* (Fig 5 A-C) and *Smo^LoF/+^* mice (which were indistinguishable from wildtype), with multiple, large ectopic signals apparent around the knee (Fig 5 A-F) and ankle joints (Fig 5 H-M). Volumetric analysis revealed a significant increase in the volume of *Ubr5^mt^+Smo^LoF^* femoral-associated ectopic bodies compared to *Ubr5^mt^* alone (Fig. 5G) and the ankles harboured a 20-fold increase in the volume of ectopic signals (Fig 5O). In agreement, histological analysis of the *Ubr5^mt^+Smo^LoF^* joints revealed an enhanced phenotype to that described in *Ubr5^mt^* (Figs. 2 & 3). *Ubr5^mt^+Smo^LoF^* synovium harboured large ectopic tissue masses (Fig 6A) with extensive vascularisation (Fig 6B) and chondrocytes lining the surface (Fig 6C) with deeper calcified cartilage and vascularization (Fig 6D). Sagittal sectioning through the ankle revealed large ectopic structures within the superficial digital flexor tendon (Fig 6E), consisting of bone and cartilaginous tissue (Fig 6F, H), and at the tendon interface (Fig 6G). Large swathes of chondrocytes were present within the superficial digital flexor and AT that coincided with an absence of tenocytes (Fig 6I, J), as previously reported in the *Ubr5^mt^* (Fig. 2). In addition, the AC in *Ubr5^mt^+Smo^LoF^* knee joints exhibited extensive loss over both tibial and femoral surfaces at this young age (Fig. 6M, N), while *Ubr5^mt^* knee joints exhibited only tears within the AC (Fig 6 K, L, quantification in O). Importantly, the loss of a single copy of *Smo* alone *(Prx1-Cre;Smo^LoF/+^)* resulted in no structural or AC damage (Fig 6P).

**Figure 5.**
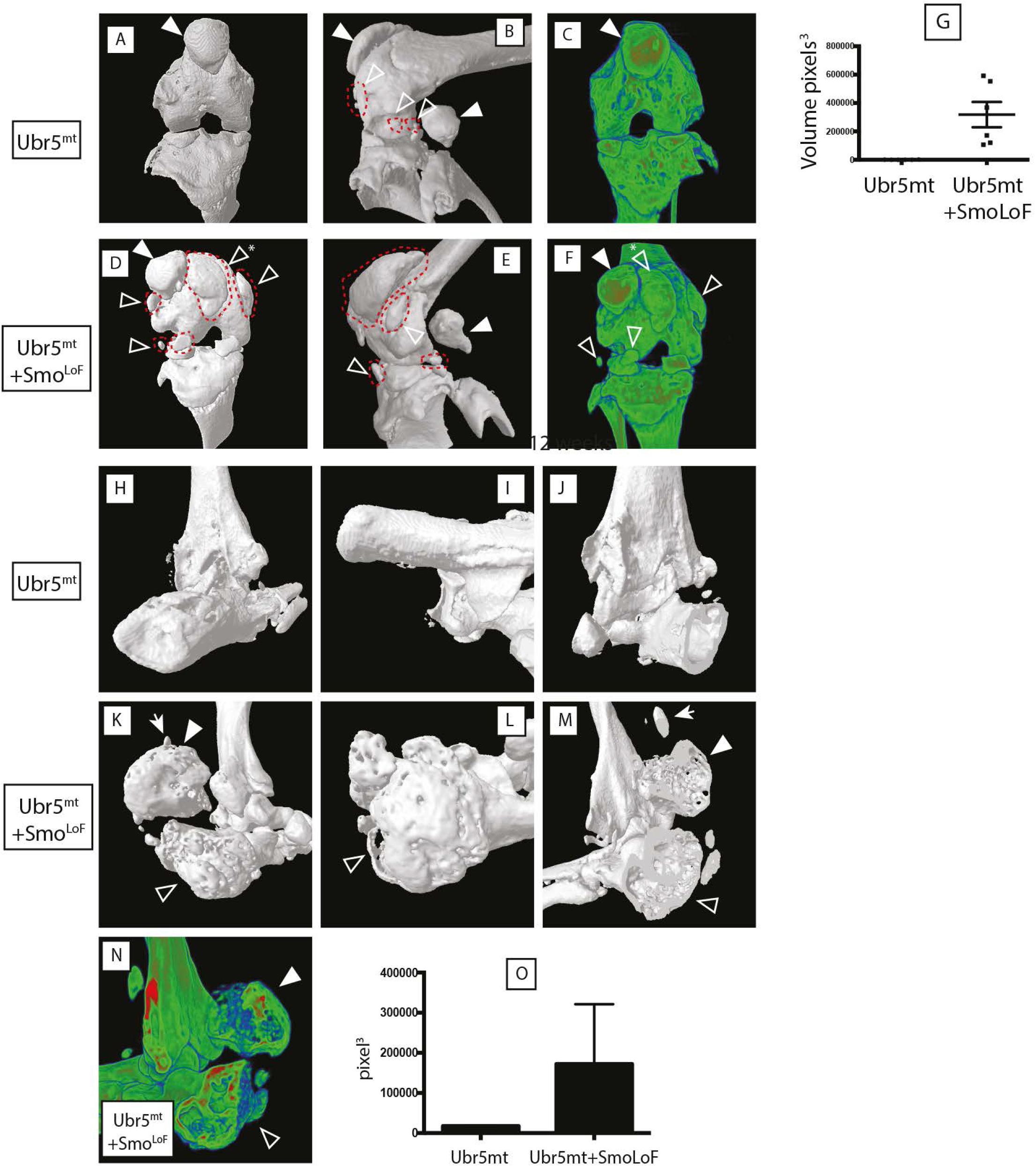
*Smo^LoF^* enhances the *Ubr5^mt^* ECCO phenotype. Analysis of 12-week-old knee (A-G) and ankle (H-O) joints by μCT-based 3D models. (A-C) *Ubr5^mt^* and (D-F) *Ubr5^mt^+Smo^LoF^* knee joints revealed ectopic structures marked by red dashed lines and open arrowheads. Sesamoid bones indicated by closed arrowhead. Asterisk marks an ectopic structure displacing the patella. All images of surface rendered 3D models, except for (C,F) that are volume rendered. (G) Volumetric analysis of ectopic structures revealed *Ubr5^mt^+Smo^LoF^* exhibited a dramatic increase in total ectopic volume over *Ubr5^mt^* alone. Mean and s.e.m indicated. n = six knees from three animals for each genotype. t-test. p = 0.0002. (H-J) *Ubr5^mt^* ankle joints exhibited a few small ectopic signals. (K-N) *Ubr5^mt^+Smo^LoF^* ankles joints exhibited large (closed arrowhead) and small (arrow) ectopic signals in addition to an abnormal and enlarged calcaneus (open arrowhead). (N) Optical cross sections through volume-rendered model revealed the internal structure and X-ray densities of the (open arrowhead) calcaneus and (closed arrowhead) ectopic structure. (O) Volumetric quantification of ectopic structures in the indicated genotypes. n = five animals per genotype. t test, p value = 0.0293. Mean and s.e.m. indicated.

**Figure 6.**
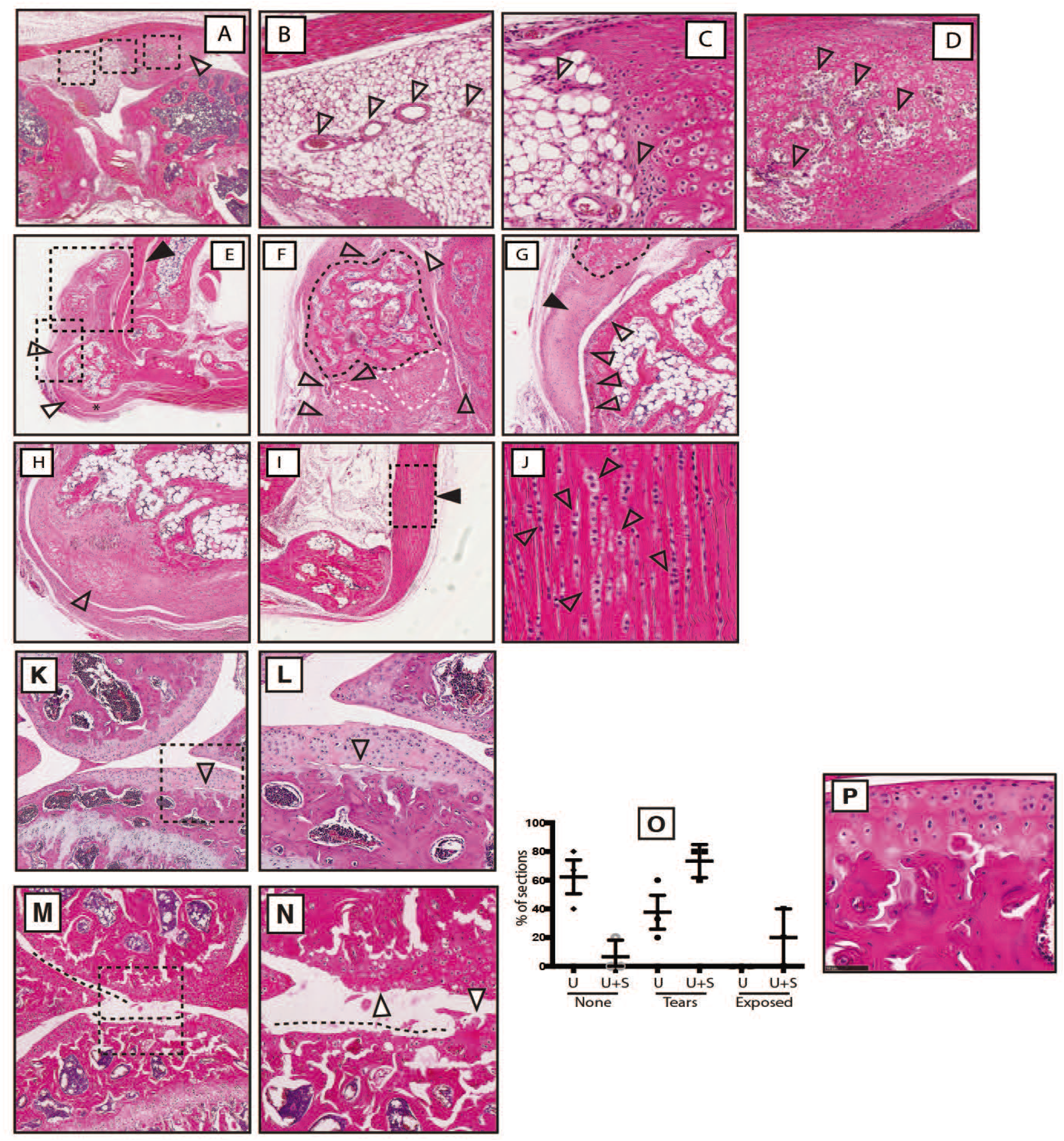
*Smo^LoF^* enhances the *Ubr5^mt^* AC phenotype. (A) *Ubr5^mt^*+ *Smo^LoF^* synovium exhibited large ectopic tissue deep to the patella and adjacent to the femur (open arrowhead). Three black dashed boxes, from left to right, are enlarged in (B), (C) and (D), respectively. (B) Sub-intimal synovial layer abutting the ectopic tissue was highly vascularized (arrowheads). (C) The region interfacing with the ectopic tissue harboured plump spindle-like cells and chondroid-like cells (arrowheads). (D) The core of the ectopic tissue resembled calcified cartilage undergoing endochondral ossification and harboured vascularized cavities (arrowheads). (E) *Ubr5^mt^+Smo^LoF^* ankle exhibited a large ectopic structure (black arrowhead) and abnormal superficial digital flexor tendon (open arrowhead) and calcaneus (asterisk). The upper and lower dashed boxed region are enlarged in (F) and (G), respectively. (F) The ectopic mass contained bone-like (black dashed lines) and cartilaginous tissues (white dashed lines) surrounded by extensively vascularised synovium (open arrowheads). (G) The presumed superficial digital flexor tendon attached to the ectopic bone (black arrowhead), harboured chondrocytes and resembled cartilage (encircled by black dash line). The adjacent periosteum of the calcaneus was highly vascularised (open arrowheads). (H) Cartilaginous thickening of the outer calcaneus. (I) The AT was thickened. The dashed box enlarged in (J) shows columns of chondrocytes (open arrowheads). (K-N) Analysis of 12-week-old knee joints of *Ubr5^mt^* and *Ubr5^mt^+Smo^Lof^* by H&E stained histological sections of the lateral condyles. (K, L) *Ubr5^mt^* exhibited tears in the AC (open arrowhead) and (M, N) *Ubr5^mt^+Smo^LoF^* exhibited extensive loss of AC (dashed lines) and damaged apical surfaces (arrowhead). Dashed boxes in (K) and (M) indicate the enlarged regions in (L) and (N), respectively. (O) Percentage of sections bearing no damage (‘None’), tears (‘Tears’) or exposed calcified cartilage (Exposed), revealed a significance difference between the genotypes. Mean and s.e.m indicated. Chi-square test on pooled slide counts. p = 0.0027. (P’) *Smo^LoF^* AC showed no signs of AC damage.

### *Ubr5* suppresses canonical HH signalling and PKA activity

A functional link between UBR5 activity and HH signalling was further examined in 6-week old *Ubr5^mt^* mice. At this age ectopic structures were not detectable (Fig. 4), thereby increasing the likelihood of detecting potential causative changes in expression patterns. Immunohistochemistry on *Ubr5^mt^* knee intimal (Fig 7 A-F) and subintimal synovium (Fig 7 G – L) revealed increased *Gli1* expression in comparison to *Prx1-Cre* control animals (Fig 7 B, E and H, K; respectively), indicative of increased canonical HH signalling. qRT-PCR analysis also confirmed increased expression of markers of canonical HH signalling in RNA from isolated synovium *(Gli1* and *Ptc1)* (Fig. 7M). Additionally, intimal and sub-intimal *Ubr5^mt^* synovium exhibited increased phosphorylated PKA substrate (PPS) staining suggesting decreased Gi proteins activation, characteristic of non-canonical HH signalling (Fig. 7 C, F, and I, L). Consistent with the observations in the synovium, *Ubr5^mt^* AC exhibited markers of increased canonical (Fig 8 A-D) and decreased non-canonical HH signalling (Fig 8 E, F). Although little change for PTCH1 was detected (Fig 8G) there was significant differences for Gli1 expression and PKA substrate staining (Fig 8 G-I).

**Figure 7.**
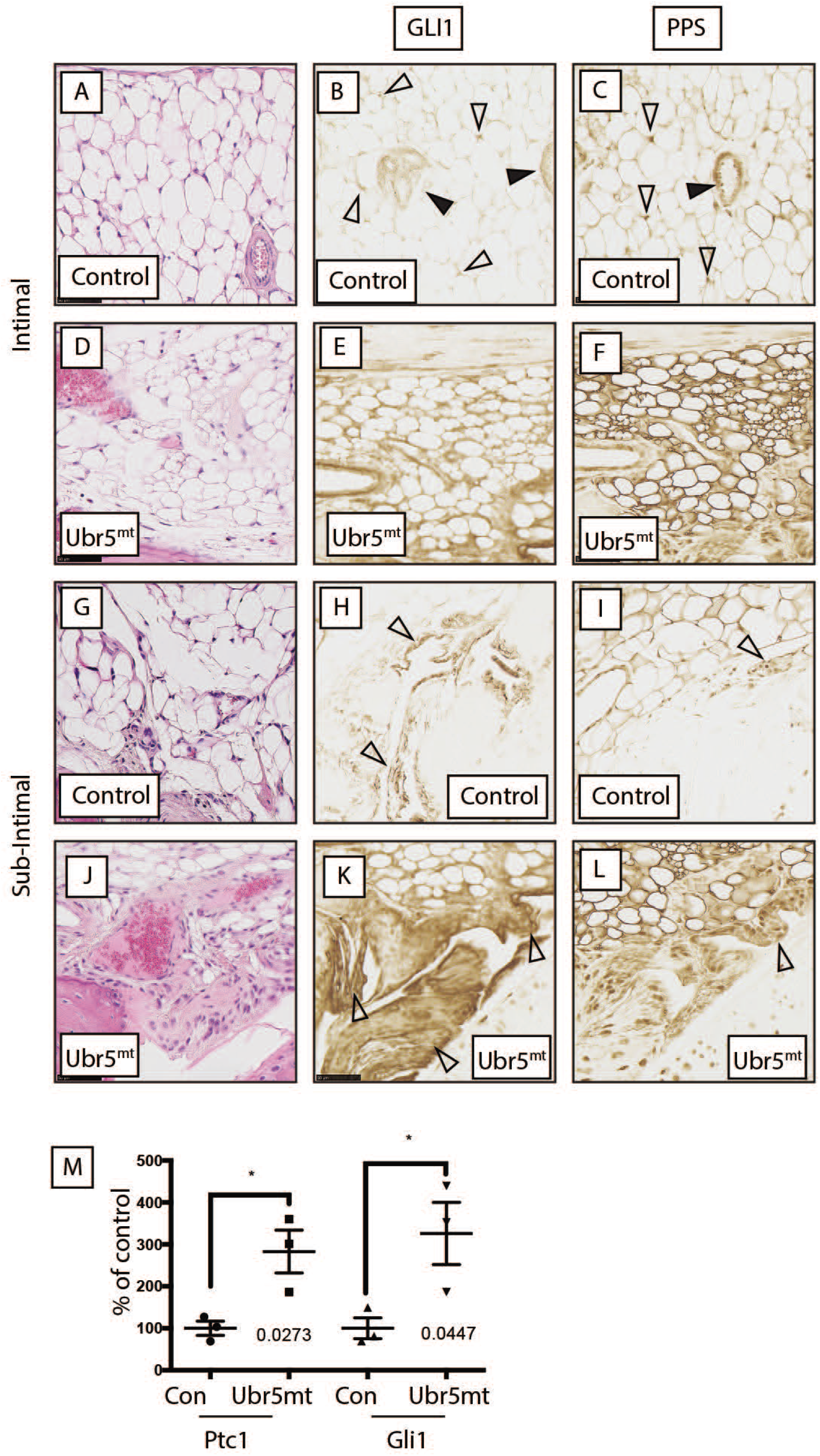
*Ubr5^mt^* synovium exhibits markers of increased canonical and decreased non-canonical HH signalling. Immunohistochemical localization of markers of canonical and non-canonical HH signalling (A-L) six-week-old sagittal sections of *Prx1-Cre* control (Con) and *Ubr5^mt^* animals. In general, control (A-C) intimal and (G-I) subintimal layers exhibited weaker GLI1 and PPS staining than in comparable *Ubr5^mt^* sections (D-F and J-L, respectively). (B) GLI1 staining in the control intimal layer was located to the vasculature (closed arrowheads) and some adipocytes (open arrowheads). (C) PPS staining was in the vasculature (closed arrowheads) and within adipocytes (open arrowheads). (E) GLI1 and (F) PPS staining were throughout the subintimal layer. (H) GLI1 staining of control synoviocytes within the subintimal layer (arrowhead). (I) PPS staining in subintimal layer synoviocytes (arrowhead). (K) GLI1 and (L) PPS staining were strongly expressed within hyperplastic and thickened synovial sub-intimal layer. Synoviocytes exhibited robust nuclear and cytoplasmic staining for GLI1 (K) and PPS (L). (M) qRT-PCR on synovium RNA for expression of canonical HH pathway expression markers *Ptch1* and *Gli1.* Graph indicates mean and s.e.m. n = three animals. t-test. *Ptch1* p = 0.0273 and *Gli1* p = 0.0477.

**Figure 8.**
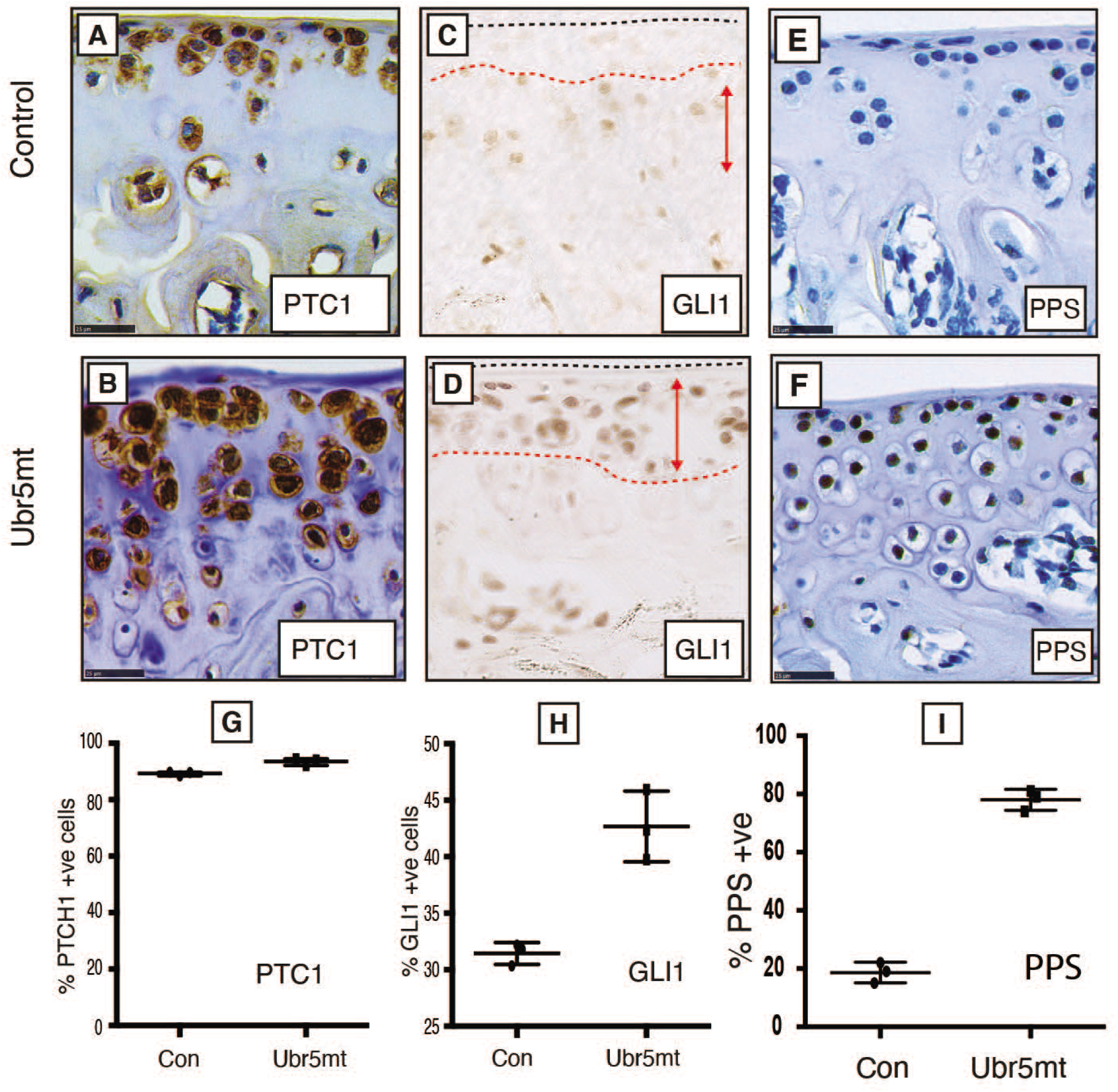
Impaired Ubr5 function results in increased canonical and decreased non-canonical HH signalling. Immunohistochemical analysis of six-week-old control and *Ubr5^mt^* tibial AC examined for markers of canonical HH pathway activity. Relative to (A, C) control, (B, D) *Ubr5^mt^* AC displayed increased staining intensities for PTCH1 (A, B) and GLI1 (C, D) with GLI1 exhibiting expanded expression domains (D, double-headed arrows). (E, F) Staining for PKA phosphorylated substrates (PPS) revealed (Q) *Ubr5^mt^* AC exhibited increased numbers of robust staining cells. The number of expressing cells is quantified in (G-I). Quantification confirmed *Ubr5^mt^* AC to harbour increased numbers of positive cells for all antigens except PTCH1. Graphs represent the percentage of positive cells, regardless of staining intensity, with the mean and s.e.m indicated. n=3 biological replicates. Chi-square test on pooled cells count data. p = <0.0008, except PTCH1 which was not significant.

### UBR5^mt^ AC and damaged human AC exhibits both aberrant expression of markers of chondrogenesis and HH signalling

As seen in murine *Ubr5^mt^* AC, osteoarthritic AC from patients also exhibits markers of increased canonical HH signalling [32]. We next addressed (i) UBR5 expression and (ii) markers of decreased non-canonical HH signalling (PPS) in human AC. Graded samples from (OA) patients (S4 Fig A-C) undergoing total joint replacement were assessed for UBR5 expression (S4 Fig E, G, I) and PKA activity (PPS in S4 Fig D, F, H). As in the murine model, PPS IHC staining increased (S4 Fig J), and hUBR5 staining decreased (S4 Fig K) with decreasing AC health. Observations of changes in markers consistent with increased canonical and decreased non-canonical HH signalling in *Ubr5^mt^* synovium and AC were echoed in human OA samples.

To further delineate whether mammalian *Ubr5* could influence markers of canonical and non-canonical HH signalling, murine NIH3T3 cells were engineered to either exhibit increased (cDNA overexpression) or decreased (shRNA knock-down) *Ubr5* expression. Cells were then transfected with constructs encoding (i) *Shh*, (ii) constitutively active *Smo* mutant *(Smo-M2)* [35] or (iii) *Gli1.* Canonical pathway activity was measured using a *Gli*-responsive luciferase reporter assay. While perturbation of *Ubr5* expression had no effect on Shh- or Smo-M2-mediated signalling (Fig. 9A and 9B), *Ubr5* overexpression caused a significant reduction (Fig. 9A, P<0.001), and *Ubr5* shRNA-mediated knockdown caused a significant increase (Fig. 9B, P<0.05), in Gli1-mediated luciferase activity. However, *Ubr5*-overexpression did not perturb the expression level of endogenous or exogenous GLI1 protein (Fig. 9C), excluding a role for UBR5-mediated degradation. Therefore, UBR5 appeared to only suppress canonical HH signalling associated with overexpression of GLI1.

**Figure 9.**
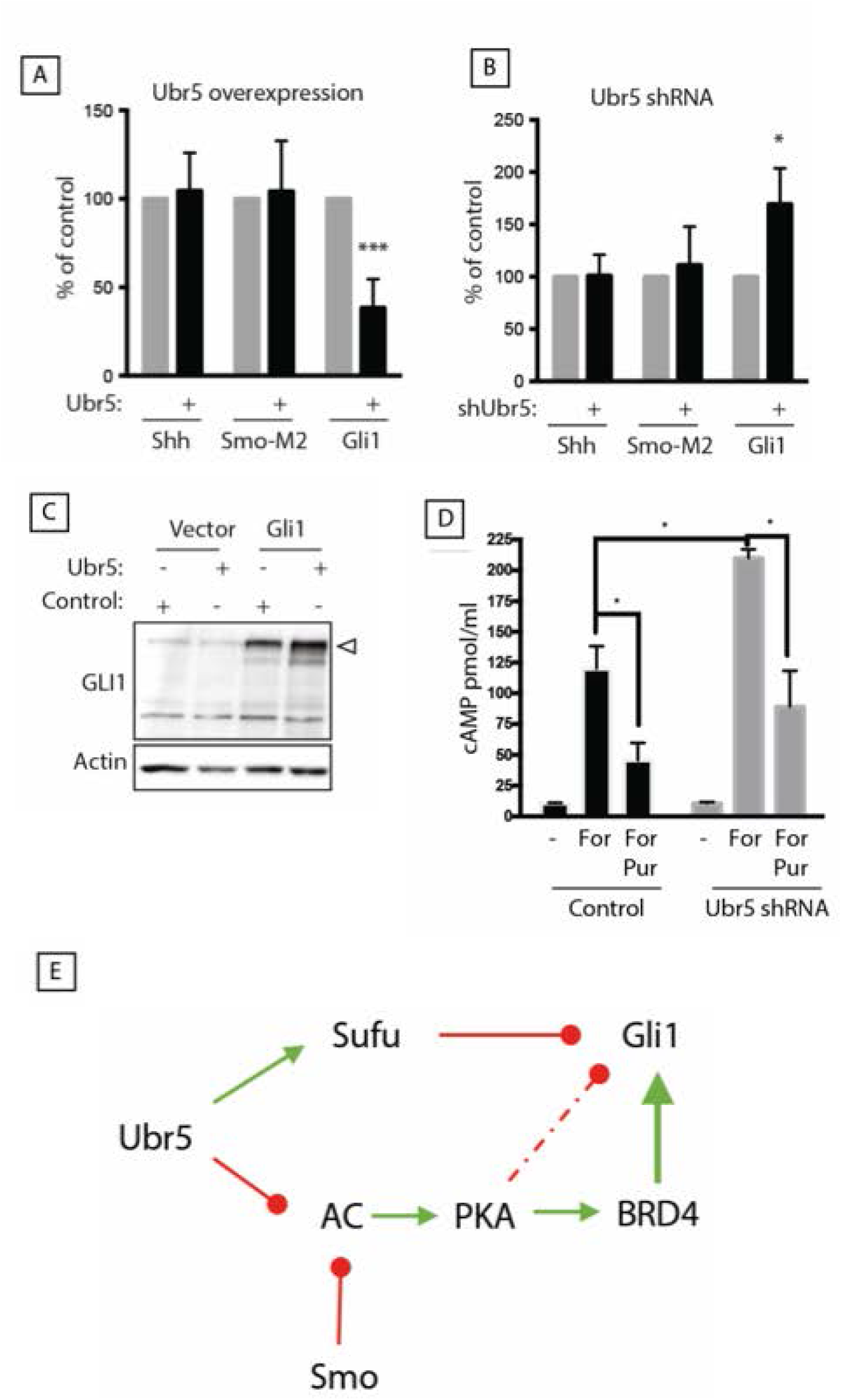
Ubr5 functions as a negative regulator of HH signalling ex vivo. Analysis of HH pathway activity in murine NIH3T3 cells in response to modulation of Ubr5 expression. (A) Cells were transfected with empty pN21 vector (grey bars) or pN21-*Ubr5* (black bars) together with plasmids encoding *Shh, Smo-M2* or *Gli1* and *8xGLI-Firefly* and *pTK-Renilla* luciferase reporters in growth medium (DMEM with 10% FBS). After 24 h, serum was reduced to 0.5% and the Firefly/Renilla luciferase activity was measured 48 h later. Bars represent mean +/− s.e.m. of n = 3 independent experiments. (B) A similar GLI-luciferase assay was carried out in NIH3T3 cells stably expressing *Ubr5* shRNA (black bars) or *scrambled* shRNA (grey bars) Bars represent mean +/− s.e.m. of n = 3 independent experiments. (C) NIH3T3 cells were cotransfected with pN21-*Ubr5* (Ubr5) or empty vector (Control) and *Gli1-myc* or empty pcDNA3.1, followed by Western blot analysis of Gli1 expression (arrowhead) using □-actin as loading control. (D) Stable knockdown of *Ubr5* impaired readouts of non-canonical HH signalling. Production of cAMP by control *scrambled* shRNA (black bars) or *Ubr5* shRNA stable cells (grey bars) following acute treatment with the adenylate cyclase activator forskolin (For) or forskolin plus the SMO agonist purmorphamine (For/Pur), compared to DMSO vehicle as control (-). Forskolin-stimulated cAMP production in *Ubr5* shRNA cells was significantly elevated compared to control cells (p = 0.0368; *t*-test). Purmorphamine suppressed forskolin-mediated cAMP production in both scramble control (p = 0.0318; *t*-test) and *Ubr5* (p = 0.0160; *t*-test) shRNA cell lines. Graphs indicate mean and s.e.m.; n = 4 independent experiments. (E) Proposed model of UBR5 function in HH signalling: UBR5 negatively regulates canonical HH signalling downstream of SMO, hypothetically through facilitating the function of the HH negative regulator Sufu, despite simultaneously inhibiting adenylate cyclase (AC). In this context, loss of Ubr5 could increase Gli1 expression by two means: 1) impairment of Sufu negative regulation and 2) stimulation of Gli1 transcriptional activity by increasing PKA-dependent phosphorylation of BRD4. The convergence of Ubr5 and SMO to suppress adenylate cyclase activity could explain the phenotypic enhancement observed in compound mice with loss of function of Ubr5 and Smo. Green and red arrows indicate established modes of activation and repression, respectively.

We then addressed whether loss of *Ubr5* function would also affect cAMP production as a readout of Gi protein activity, an indirect marker of non-canonical HH signalling. *Ubr5* shRNA cells showed an ~2-fold increase in maximal cAMP production in response to forskolin, an adenylate cyclase agonist (Fig 9D) [33]. Moreover, simultaneous addition of forskolin and purmorphamine, a SMO agonist, lowered maximal cAMP generation, but its effect was suppressed by *Ubr5* shRNA (Fig 9 D). Together, the *in vitro* findings suggest that Ubr5 loss results in reduced stimulation of Gi proteins by Smo, leading to increased cAMP/PKA activity levels. Overall, these data supported our *in vivo* observations that *Ubr5* normally acts to suppress GLI1 activity while promoting PKA activity.

## Discussion

### *Ubr5* mutation causes musculoskeletal tissue defects

We report a role for mammalian *Ubr5* in adult skeletal homeostasis that impacts upon and genetically interacts with, components of the HH signalling pathway. These findings add to the emerging importance of the N-end rule ligases in regulating important signalling and cellular processes in human, and animal health and disease [34, 35]. Loss of the *Ubr5* gene in early limb mesenchyme resulted in postnatal defects in and around joints within the fore and hind-limb. Defects included ectopic bone and cartilage formation, and articular cartilage degradation (see summary S4 Fig 4).

Our data indicates metaplastic production of chondrocytes and/or ectopic endochondral ossification as a major component of *Ubr5^mt^*-associated ECCO. Comparison of the *Ubr5^mt^-* associated ECCO phenotype with that of human inherited HO diseases reveals some similarities and differences. Within the ECCO-prone tissues there were distinct tissue-specific responses; for example, the knee-associated synovium underwent ectopic chondrogenesis, calcification and ossification to produce bone, whereas the Achilles tendon only underwent ectopic chondrogenesis and calcification. The abnormalities of the knee-associated synovium which display heterotopic chondrogenesis are reminiscent of human benign bone tumours called osteochondromas [36] whereas, the heterotopic tissue calcification without ossification seen in the AT resembles a form of calcific tendinopathy [37]. The mouse *Ubr5* mutation, thus, provides a genetic model for the generation of these bone abnormalities and suggests that the processes of chondrogenesis, tissue calcification and ossification represent discrete, albeit interrelated, steps that when deregulated can individually, or collectively, contribute to distinct tissue pathologies.

Our findings also demonstrated an important role for Ubr5 in regulating AC homeostasis, where its loss led to dramatic cellular, extracellular and structural defects. The observed defects in HH signalling could have been causative in nature as HH signalling is intimately linked to both stem cell [22] and chondrocyte biology [10]. One of the most distinctive *Ubr5^mt^* AC defects was the tearing along the tidemark between non-calcified and calcified cartilage. This focal failure suggested the interface was prone to transverse shear forces and ‘slipping’ of one layer (i.e., noncalcified cartilage) relative to the other (i.e., calcified cartilage). Interestingly, this mode of AC shedding and the associated regions of necrosis mirrored defects observed in mammalian osteochondrosis [38, 39].

### UBR5 influences markers of canonical and non-canonical HH signalling

Based on the current dogma, we hypothesized that the *Ubr5^mt^*-associated ECCO was caused by increased HH signalling. In contrast, the introduction of *Smo^LoF^* heterozygosity into a *Ubr5^mt^* background both (i) exacerbated *Ubr5^mt^*-associated defects as well as elicited novel defects not observed by loss of *Ubr5* function alone (e.g., ECCO of the calcaneal periosteum and the superficial digital flexor tendons and increased volume and altered shape of normotopic sesamoid bones). This combined ability to influence both normotopic and heterotopic bones (S4 Fig for summary of ECCO phenotype), highlights the importance of *UBR5* ain normal and pathological skeletal tissue homeostasis. Furthermore, our genetic analysis exposed a pro-homeostatic function for SMO – and by extension HH signaling – in suppressing *Ubr5^mt^* ECCO. *In vivo* and *in vitro* observations identified a loss of *Ubr5* associated with predictors of increased (GLI1 activity) and decreased (PKA activity) canonical HH signalling. Based on the current dogma, it is difficult to reconcile increased GLI activity in the context of increased PKA activity, given that PKA phosphorylates other GLI family members, GLI2 and GLI3, targeting them for processing into transcriptional repressors [14, 40]. However, the evolving breadth of the HH pathway (Fig 9F) provides potential mechanistic explanations for this apparently paradoxical observation.

Recent evidence expanded the role of PKA to promote canonical HH signalling by promoting BRD4-mediated stimulation of GLIs transcriptional activity (Fig. 9E) [41–43]. Interestingly, HO-associated with increased HH signalling was suppressed by the BRD4 inhibitor JQ1 [44], which clearly demonstrated a role for a cAMP-PKA-BRD4-GLI1 axis in skeletal tissue homeostasis. A non-canonical role of SMO as a G protein-coupled receptor (14, 15) provides a mechanism to control PKA activity. Upon stimulation, SMO activates heterotrimeric Gi proteins, which, upon dissociation, inhibit adenylate cyclase through the Gα subunit to reduce cAMP production and PKA activation [15, 45, 46]. Therefore, SMO inhibition can lead to increased cAMP-mediated PKA activity accounting for SMO modification of the *Ubr5^mt^* phenotype, as impairment of either UBR5 or SMO leads to increased cAMP-mediated PKA activity – with their combined impairment leading to either additive or synergistic effects. Interestingly, our preliminary research (personal communicationNDGR) supports a role for UBR5 in regulating readouts of non-canonical HH signalling other than PKA (i.e.; RhoA) [16]. Although our data reveal a genetic interaction between UBR5 and an essential component of the HH signalling pathway, we cannot fully establish the underlying mechanism(s) driving *Ubr5^mt^*-associated ECCO. Future work will require developing the tools to differentiate between causative individual, or combined, contributions of aberrant canonical or non-canonical HH signalling. The addition of *Smo^LoF^* into a *Ubr5^mt^* background would have exacerbated a pre-existing imbalance between the pathway outputs to drive ECCO.

The importance of balanced canonical and non-canonical HH signalling was recently demonstrated in osteogenesis [47]. Loss of the cilia regulatory protein IFT80 resulted in impaired osteoblast differentiation and coincided with (i) decreased expression of canonical target genes and (ii) increased non-canonical activity. The authors proposed that the non-canonical HH pathway prevented, and the canonical pathway promoted, formation of osteoblasts. Due to the emerging importance of non-canonical HH signalling [12], we also propose that the combined effects on canonical and non-canonical HH signalling contributed to the observed loss of tissue homeostasis in *Ubr5^mt^* animals. Overall, our detection of *Ubr5^mt^*-associated increased canonical (GLI1 activity) and indications of decreased non-canonical HH signalling (cAMP-PKA) are in general agreement with a reported pro-osteogenic environment conducive to HO [47]. UBR5 may therefore join IFT80 [47] and DYRK1B [48] as differential regulators of canonical and non-canonical HH signalling. Our future work will involve establishing which of the various non-canonical, SMO’s GPCR-associated downstream effectors (e.g., PKA, RHOA, RAC1, PI3K etc.) [49, 50] drive ECCO.

In summary, we reveal a previously unknown role for *Ubr5* in influencing HH signalling, tissue homeostasis and preventing spontaneous ECCO. A role for UBR5 in regulating HH signalling and tissue homeostasis supports the classification of human *UBR5* as a Tier 1 human cancer susceptibility gene (Sanger Cancer Gene Consensus). We believe the *Ubr5^mt^* mouse model could assist in uncovering mechanisms that lead to disorders including characterisation of early pathological events and elucidation of pro-homeostatic mechanisms capable of promoting general bone health. In the future, manipulation of human *UBR5* and *SMO* function could potentially provide a means of preventing pathological, and promoting beneficial, chondrogenesis and ossification in both the clinic and in biomedical engineering applications.

## Materials and methods

### Human Material

Human AC was obtained from knee joint arthroplasty specimens with ethical approval from the Lothian Research Ethics Committee.

### Murine studies

Animal studies were approved by the MRC IGMM ‘Animal Care and Use Committee’ and according to the MRC ‘Responsibility in the Use of Animals for Medical Research’ (July 1993), EU Directive 2010 and UK Home Office Project License no. PPL 60/4424.

*Prx1-Cre;Ubr5^gt/gt^* experimental animals (referred to as *Ubr5^mt^)* and their respective littermate controls were generated and all experiments were conducted in accordance with the ARRIVE guidelines. Tamoxifen (0.1mg/kg body weight) in corn oil, or vehicle only, were administered i.p to six-week-old animals on two consecutive days. For X-gal staining, embryos and postnatal hind limbs were dissected, fixed in 4% formaldehyde (from paraformaldehyde (PFA)) at 4°C, washed and stained in X-Gal stain solution (XRB supplemented with 1mg/ml X-Gal) overnight [20].

### Histology

Hindlimbs were fixed in 4% formaldehyde (fromPFA)) for 72hrs at 4°C before being decalcified 0.5M ethylenediaminetetraacetic acid (EDTA) pH7.4 at 4°C. Samples were embedded in paraffin wax blocks and 5μm sagittal sections cut. For cryotome sectioning, samples were equilibrated in a 30% sucrose/phosphate buffered saline (PBS) solution at 4°C and then embedded in OCT compound (Fisher Scientific, Loughborough, UK) before 10μm sagittal sections were cut. For human material, 8×3mm blocks of AC were cut from femoral tibial condyles and fixed in neutral buffered formalin and then paraffin wax embedded. Histological staining with Von Kossa (Abcam, Cambridge, UK), toluidine blue (Sigma) and haematoxylin and eosin (Sigma) were carried out according to standard procedures. All histological scoring was carried out on the lateral tibial condyle with AC damage determined by a binary scoring system, of ‘normal’ or ‘damaged’. At least three slides separated by 25μm were analysed for each limb. For cell and immunohistochemical scoring, cell-types or positive staining cells were expressed as a percentage of the total chondrocyte count. The number of empty lacunae were expressed per mm of AC analysed.

### Immunohistochemistry

*Primary antibodies:* rabbit anti-IHH (1:200, Millipore, Billerica, US); goat anti-PTCH1 (1:50, Santa Cruz, Dallas, US); rabbit anti-GLI1 (1:50, Cell Signalling); rabbit anti-SOX9 (1:50 Santa Cruz); rabbit anti-RUNX2 (1:250, Sigma); PKA phosphorylated substrates (1:150, Cell Signalling); rabbit anti-EDD1 (HsUBR5) (1:100, Bethyl Labs, Montgomery, US). Biotinylated secondary antibodies: goat anti-rabbit and horse anti-goat (1:200, Vector Labs).

Paraffin sections were de-waxed, blocked for endogenous peroxidase and underwent antigen retrieval in 10mM sodium citrate pH6 at 80°C for 30-60 minutes. Slides were blocked with serum-free pan-species block (DAKO, Glostrup, Denmark), incubated with primary antibodies overnight at 4°C, and incubated with biotinylated secondary antibodies for 45mins at room temperature. Sections underwent streptavidin-mediated signal amplification (ELITE ABC, Vectorlabs, Burlingame, US) prior to incubation with peroxidase substrate kit DAB (Vectorlabs).

### μCT image processing

Fixed limbs were imaged at 18μm resolution using a Skycan 1076 (Bruker, USA, MA). Raw μCT image stacks was reconstructed and CTAn (Bruker) used for selecting regions of interest and acquiring 2D density maps, volumetric quantification of ectopic structures and generation of surface rendered 3D models (visualized in CTVol). For 3D density mapping of the tibial epiphysis, individual pan-, low- and high-3D density map models were combined using CTVol.

### RNA extraction and q-RT-PCR analysis

Individual joint components were micro-dissected and stored in liquid nitrogen. RNA was extracted using Trizol reagent (Life Technologies), according to manufacturer’s instructions. RNA was reverse-transcribed using QuantiTect Reverse Transcription Kit (Qiagen). The qRT-PCR was performed using LightCycler® 480 SYBR Green I Master (Roche, Germany) and target gene expression normalized to *Rpl5* and analysed using the ΔΔCT method [51].

### Plasmid constructs

The *Shh* and *SmoM2* (W593L) expression vectors were provided by P. Beachy (Stanford University, USA, CA). *mGli1* expression and the reporter vectors *8xGBS-luc* were a gift from H. Sasaki (Osaka University, Japan). *pCMV-dR8.2 dvpr* (8455) and *pCMV-VSV-G* (8454) were generated in the Weiner lab and obtained from Addgene (USA). *pRL-TK* was obtained from Promega (USA) and *pcDNA3.1+* was purchased from Invitrogen (USA). Recombinant SHH ligand was synthesized and purified as described previously [52].

The complete *Ubr5* cDNA was synthesised from murine embryonic stem cells total RNA [9] and cloned into a modified pcDNA5/FRT vector (Life Technologies) containing an amino-terminal 2×HA/2×Strep. NIH3T3 cells (American Type Culture Collection, USA) were seeded at a density of 100,000/ml and transfected after 24hr with *pcDNA3.1* alone, *UBR5* and *pcDNA3.1, Gli1* and *pcDNA3.1,* or Gli1 and *Ubr5* using FuGENE6 (Roche). After 48hrs, the medium was replaced by DMEM/0.5% FCS, and cells were lysed 24 hrs later in Laemmli buffer. Whole cell lysate was separated on a 6% SDS-PAGE and transferred onto nitrocellulose membranes. Membranes were blocked in 5% non-fat milk, incubated with primary antibodies overnight at 4°C at 1:1,000 dilution for GLI1 (Cell Signalling) or 1:10,000 dilution for β-actin (Sigma). Secondary HRP-conjugated-anti-mouse antibody was applied at a 1:2,000 dilution for 1hr at room temperature. The membranes were developed using the Clarity western ECL substrate (BioRad, USA, CA).

### Retrovirus production and stable *Ubr5* silencing

Previously validated shRNA-encoding oligos targeting murine *Ubr5* and or a scrambled sequence were cloned into *pLKO.1-puro* (Sigma). sh*UBR5* and *shScrambled*-*pLKO.1-puro* were co-transfected with *pCMV-VSV-G* and *pCMV-dR8.2 dvpr* plasmids into HEK 293T cells using TransIT293 reagent (Mirus Bio LLC, USA). To generate stable silenced *shUBR5* cells, NIH3T3 cells were seeded at 120,000 cells/ml and infected with 0.5 ml *shScramble* or sh*UBR5* retroviral supernatant in the presence of 8 mg/ml polybrene (Sigma). The media was changed after 24 hrs and cells were selected with 2 mg/ml puromycin 48 hrs post-infection.

### *Gli*-luciferase assay

NIH3T3 cells were seeded, and after reaching 70% confluence transfected with *pcDNA3.1, Shh, SmoM2,* or *Gli1* together with *Gli*-luciferase and *Renilla luciferase* reporter plasmids with or without *pcDNA5-HA-Strep-Ubr5*, using FuGENE 6 transfection reagent (Roche) according to the manufacturer’s protocol. For *Ubr5* knockdown studies, stable *shScramble* and *shUbr5* NIH3T3 cells were transfected with *pcDNA3.1*, *Shh*, *SmoM2*, or *Gli1*, together with *Gli*-luciferase and Renilla luciferase reporter plasmids. In both cases, after the cells reached 100% confluency, the medium was replaced with DMEM/0.5% FCS. After 24 hrs, Firefly and Renilla luciferase activities were determined with the Dual Luciferase Reporter Assay System (Promega).

### cAMP assay

Control (scrambled) or knock down *(Ubr5* shRNA) NIH3T3 cells were seeded at 130,000 cells/ml, serum starved overnight, and stimulated with 10μM forskolin (FORSK) for 5min. Cells were pre-incubated with 5μM purmorphamine for 10min before addition of FORSK. Cells were processed according to Parameter cAMP Enzyme Immune Assay (R&D Systems) instructions.

### Statistical analysis

Data analysis and statistics was performed using PRISM software (GraphPad, La Jolla, US). Count data was analysed using a contingency table and either two-sided Chi square or Fisher’s exact tests according to count size. Continuous data was analysed using unpaired, two-tailed Students t-tests. The level of significance for all tests was set at p=<0.05.

## Acknowledgements

We would like to thank Lorraine Rose and Rob v’ant Hof for preliminary μCT scanning and the SURF and IGMM Histology facilities for their services. We also thank the BRF for expert technical assistance.

## Funding

MD is supported by a University of Edinburgh Chancellor’s Fellowship and funding from a Carnegie Research Incentive Grant (70356); BH and SP by a MRC core award to the MRC HGU; and KS by the MRC (MR/R022240/1). CF supported by BBSRC through an Institute Strategic Programme Grant Funding (BB/J004316/1).

## Supplementary Information

**Supplementary Figure 1.**
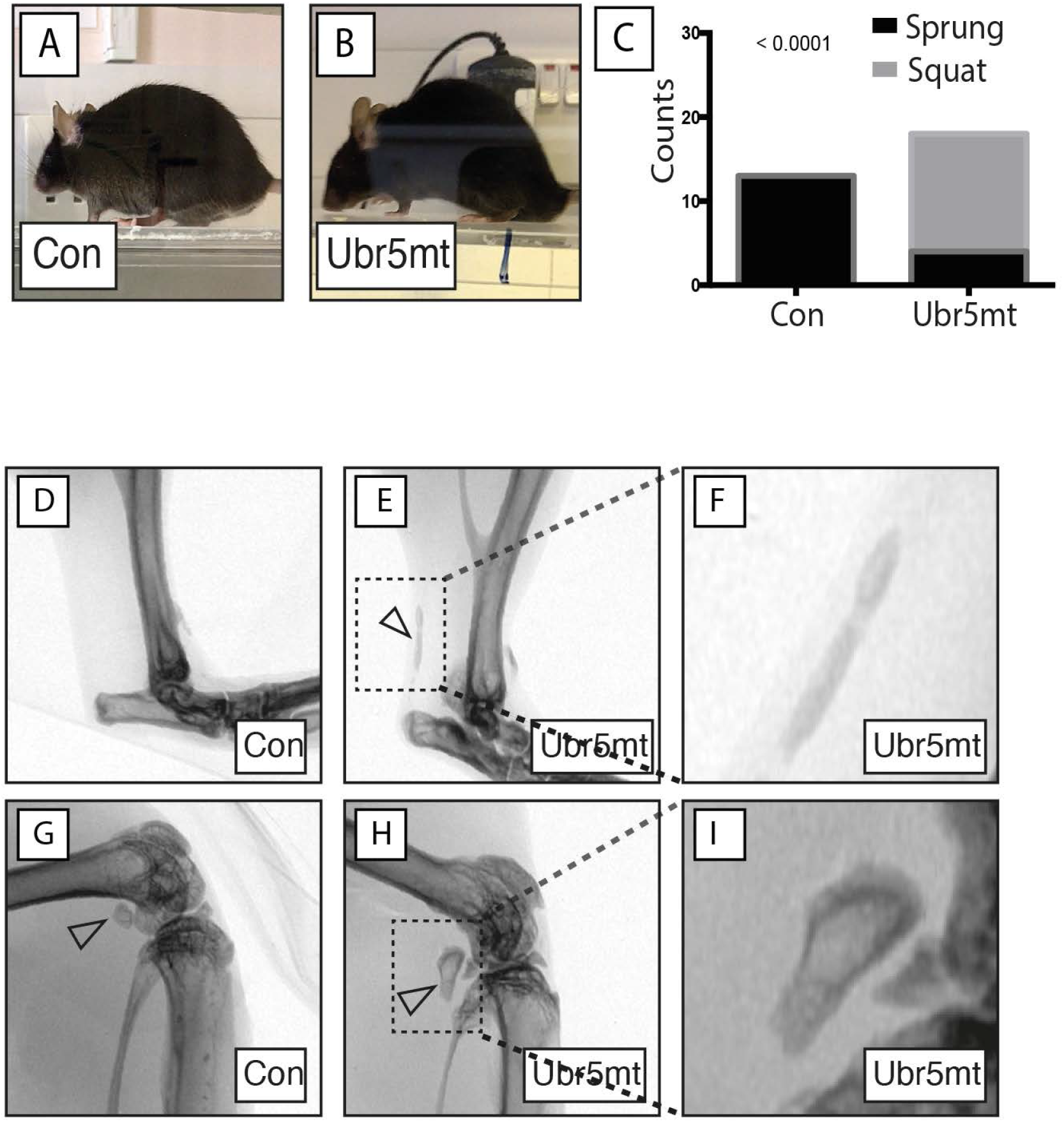
*Ubr5^mt^* mice exhibit gait abnormalities and ectopic X-ray-dense signals. 24-week-old control or *Ubr5^mt^* mice were assessed for (A-C) behavioral analysis. (A,B) Mice were videoed while walking along a boxed runway and their static positioning recorded as either ‘sprung’ (with their posterior not in contact with the floor), or ‘squat’ (with their posterior resting on the floor). (C) Graph showing counts of animal behavior. n = six and eight male and females for control and Ubr5mt genotypes, respectively. Fisher’s exact test, p value = <0.0001. (D) Control ankles and (E) *Ubr5^mt^* ankles which exhibited ventrally- and dorsally located isolated signals. (F) Dashed box region enlarged. (G) Control knee joints exhibited the fabella, a dorsally-located sesamoid bone (closed arrowhead). (H) *Ubr5^mt^* knee joints exhibited a misshapen fabella (closed arrowhead), with the dashed boxed region being enlarged (I). n = eight males and eight females.

**Supplementary Figure 2.**
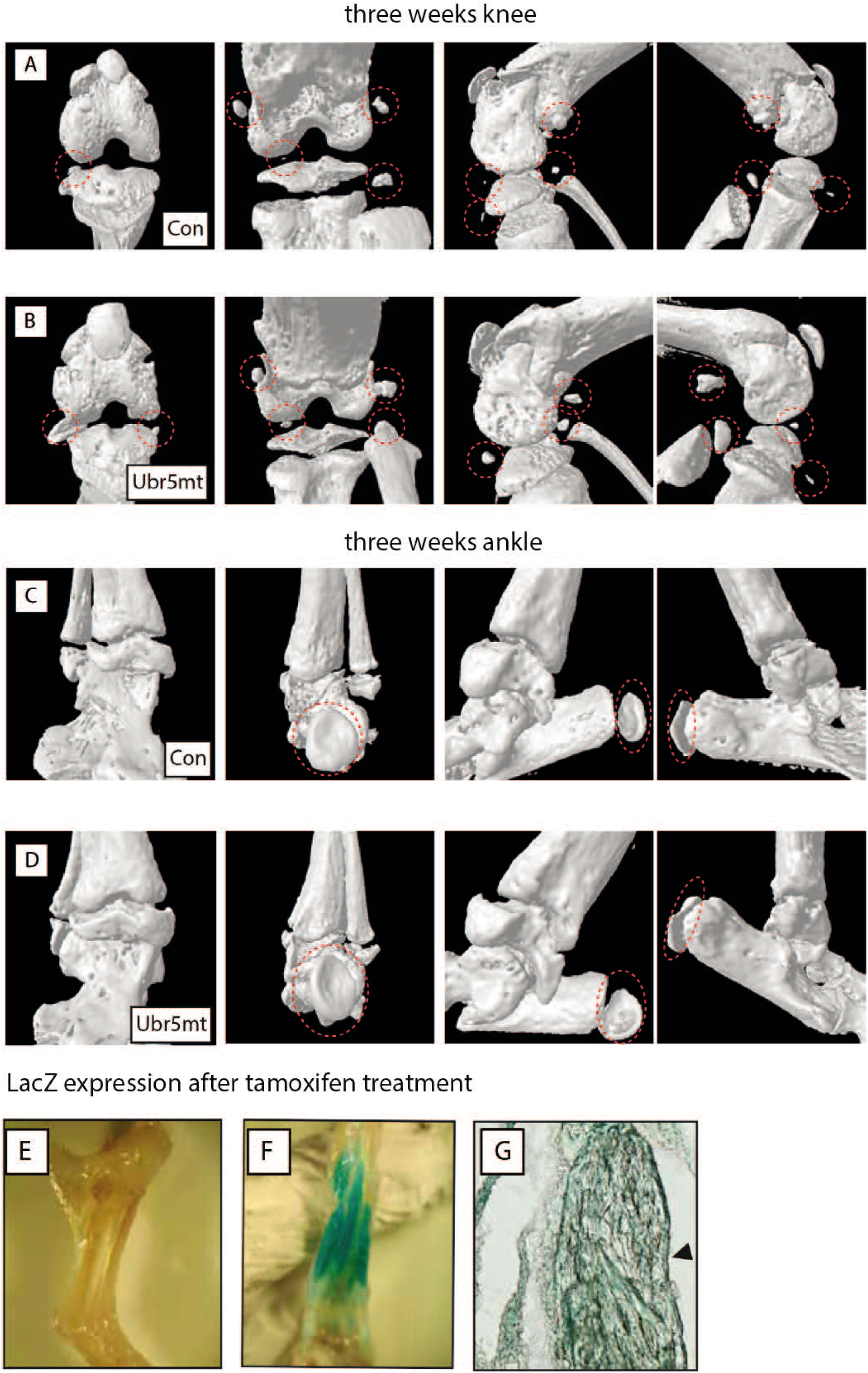
Ectopic structures are not detected in three-week-old control or *Ubr5^mt^* ankle or knee joints and require postnatal expression of *Ubr5.* (A-D) Different views of surface rendered 3D models of three-week-old control and *Ubr5^mt^* (A,B, respectively) knee and (C,D, respectively) ankle joints. (A-D) From left to right panels: ventral, dorsal, medial and lateral views. Both control and *Ubr5^mt^* joints exhibit either a normal array of sesamoid bones, developing epiphysis and calcifying menisci. (E-G) Analysis of 18-week-old tamoxifen-treated *pCAGG-Cre* control and pCAGG-*Ubr5^mt^* ankle joints. (E,F) Whole mount β-Gal staining of (E) control and (F) *pCAGG-Ubr5^mt^* ankle joints reveals β-gal expression in muscles and associate tendons. Sagittal section of ankle joint (G) showing an ectopic structure associated with the AT midbody (closed arrowhead) stained positive for *Ubr5/UBR5* expression

**Supplementary Figure 3.**
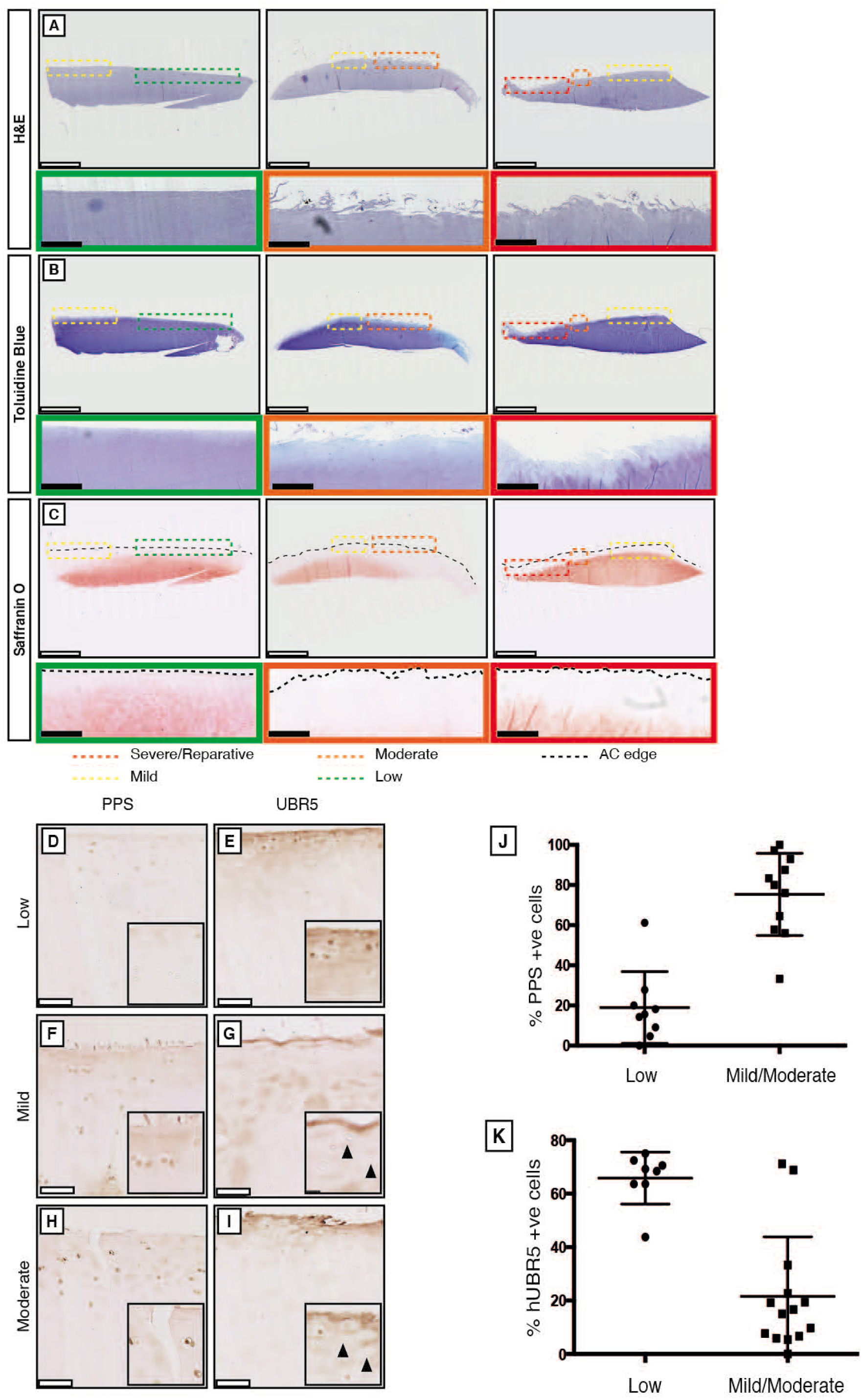
UBR5 and PPS levels correlate with human AC damage. (A-C) Examples of human OA patient material stained with (A) haematoxylin and eosin (H&E), (B) toluidine blue or (C) safranin O revealed intra- and inter-sample variation in AC defects. Coloured boxes indicate regions of the varying OA severity (please see figure key). (A-C) Colour-coded, magnified dashed boxes in upper panels are shown in more detail in the colour-coded lower panels (thick outlines). Moderate-scored regions (orange) exhibited extensive surface fibrillation and reduced toluidine blue and safranin O staining in comparison to low-scored regions (green). Severe-scored regions (red) exhibited loss of safranin O staining and apical-basal clefts in the AC surface. (C) The dashed black lines indicate the apical edge of the AC. (D-K) Human AC samples graded as low, mild or moderately damaged were analysed for (D,F,H) PKA activity (PPS) and (E,G,I) UBR5 expression. Graphs of percentage of (J) PPS and (K) UBR5 positive cells for low and combined values for mild and moderate AC grades. Mean and s.e.m indicated. n = six biological replicates. Fishers exact test on pooled cell count data. p= <0.0001 for both.

**Supplementary Figure 4.**
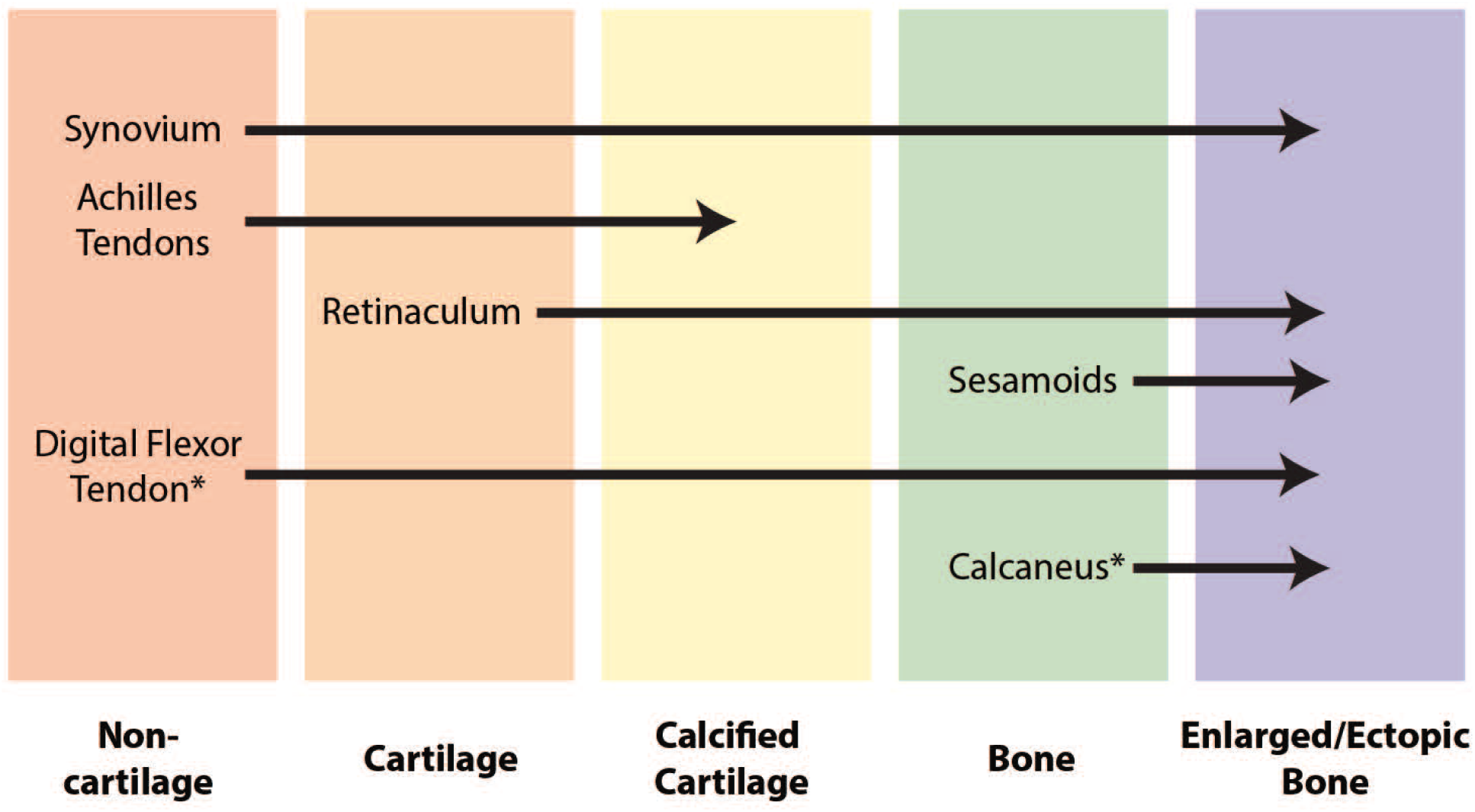
Spectrum of *Ubr5^mt^* and *Ubr5^mt^+Smo^LoF^* associated tissuespecific metaplastic responses. Overview of the metaplastic events of various *Ubr5^mt^* tissues. Red = non-cartilaginous tissues (Synovium, AT and Superficial Digital Flexor tendon); Orange = cartilaginous tissues (retinaculum); Yellow = calcified cartilage; Green = normotopic bone; Blue = heterotopic or enlarged normotopic bone. Arrows indicate the direction of metaplasia, with the arrowhead indicating the tissue type in 24-week-old *Ubr5^mt^* and/or *Ubr5^mt^+Smo^LoF^* animals. Metaplastic tissue events unique to *Ubr5^mt^+Smo^LoF^* are indicated by an asterisk.

